# Multiple Intrinsic Membrane Properties are Modulated in a Switch from Single to Dual-Network Activity

**DOI:** 10.1101/2022.08.01.502404

**Authors:** Ryan R. Snyder, Dawn M. Blitz

## Abstract

Neural network flexibility extends to changes in neuronal participation between networks. This neuronal switching can include neurons moving between single- and dual-network activity. We previously identified an example in which bursting at a second frequency occurs due to modulation of intrinsic membrane properties instead of synaptic recruitment into a second network. However, the intrinsic properties that are modulated were not determined. Here, we use small networks in the Jonah crab (*Cancer borealis*) stomatogastric nervous system (STNS) to examine modulation of intrinsic properties underlying neuropeptide- (Gly^1^-SIFamide) elicited neuronal switching. The LPG neuron switches from exclusive participation in the fast pyloric (∼1 Hz) network, due to electrical coupling, to dual-network activity which includes periodic escapes from the fast rhythm via intrinsically-generated oscillations at the slower gastric mill network frequency (∼0.1 Hz). We isolated LPG from both networks using pharmacology and hyperpolarizing current injection. Gly^1^-SIFamide increased LPG intrinsic excitability and rebound from inhibition, and decreased spike frequency adaptation, which can all contribute to intrinsic bursting. Using ion substitution and channel blockers, we found that a hyperpolarization-activated current, a persistent sodium current, and a calcium or calcium-related current(s) appear to be primary contributors to Gly^1^-SIFamide-elicited LPG intrinsic bursting. However, this intrinsic bursting was more sensitive to blocking currents when LPG received rhythmic electrical coupling input from the fast network than in the isolated condition. Overall, a switch from single- to dual-network activity can involve modulation of multiple intrinsic properties, while synaptic input from a second network can shape the contributions of these properties.

**New and Noteworthy:** Neuropeptide-elicited intrinsic bursting was recently determined to switch a neuron from single to dual-network participation. Here we identified multiple intrinsic properties modulated in the dual-network state and candidate ion channels underlying the intrinsic bursting. Bursting at the second network frequency was more sensitive to blocking currents in the dual-network state than when neurons were synaptically isolated from their home network. Thus, synaptic input can shape the contributions of modulated intrinsic properties underlying dual-network activity.

## Introduction

Flexibility of rhythmic networks enables adaptation to changing internal and external environmental demands. This network plasticity results in multiple versions of rhythmic locomotor, respiratory and chewing behaviors, and different oscillatory patterns contributing to sleep, memory formation, and other cognitive processing (Barlow 2009; Cropper et al. 2017; Dickinson 1995; Rangel et al. 2016; Roopun et al. 2008; Stickford and Stickford 2014). Network flexibility extends to neuronal switching, in which neurons change their participation between networks, including switching between single- and dual-frequency oscillations (Fahoum and Blitz 2021; Hooper and Moulins 1990; Meyrand et al. 1994; Steriade et al. 1993; Tryba et al. 2008; Weimann et al. 1991).

Neuronal switching contributes to multifunctional roles for neurons and muscles in a number of behaviors (Bouret and Sara 2005; Briggman and Kristan 2008; Dickinson 1995). For instance, the same neurons participate in combinations of coughing, gasping, and breathing in rodents and cats, and in multiple feeding-related behaviors in molluscs and crustaceans (Cropper et al. 2017; Lieske et al. 2000; Meyrand et al. 1994; Oku et al. 1994; Shannon et al. 1998; Weimann et al. 1991). Bifunctional muscles similarly switch their participation between different behaviors, such as abdominal muscles in bats that contribute to respiration and the generation of echolocation signals (Lancaster et al. 1995). In other cases, rhythmic networks and behaviors are co-active, and coordinating their activity via switching neurons is functionally important (Stickford and Stickford 2014; Wei et al. 2022). Disrupted coordination can lead to behavioral deficits such as aspirating food when swallowing (Barlow 2009; Yagi et al. 2017).

Despite the importance of coordination between networks, it is not a static parameter. In crustacean feeding networks, temperature and oxygen levels alter coordination between networks (Clemens et al. 1998; Stein and Harzsch 2021), and locomotor speed alters coupling ratios between human locomotion and respiration (Saunders et al. 2004). Thus, understanding the mechanisms controlling neuronal switching is important to understand function and dysfunction of multiple related networks.

Although neuronal switching is thought to be prevalent in many regions of vertebrate and invertebrate nervous systems, there are few instances of identified cellular-level mechanisms regulating neuronal participation between networks. In smaller, invertebrate systems the predominant mechanism identified for recruiting a neuron into an additional network, or assembling a novel network from multiple discrete networks, is modulation of synaptic strength (Gutierrez et al. 2013; Hooper and Moulins 1990; Meyrand et al. 1994). In addition to synaptic properties, intrinsic membrane properties controlling neuronal excitability are also important targets of neuromodulators that alter network output (Harris-Warrick 2011; Marder and Thirumalai 2002; Nadim and Bucher 2014). For instance, modulation of intrinsic membrane properties can “release” neurons from a network (Faumont et al. 2005; Hooper and Moulins 1990; Tryba et al. 2008). More recently, we found that modulation of membrane properties can also recruit a neuron into dual-network activity (Fahoum and Blitz 2021), as suggested in computational studies (Drion et al. 2019; Gutierrez et al. 2013). Large, diffuse networks can make it difficult to examine the cellular-level mechanisms of neuronal switching, and thus most mechanistic information comes from studies in smaller, well-defined neural networks (Bouret and Sara 2005; Briggman and Kristan 2008; Dickinson 1995).

Feeding-related networks in the STNS of the crab, *Cancer borealis*, provide an exceptional model for examining network plasticity, including neuronal switching. The stomatogastric ganglion (STG) contains approximately 30 neurons comprising two rhythmic networks; the fast pyloric (∼1 Hz; food filtering) and slow gastric mill (∼0.1 Hz; food chewing) networks (Daur et al. 2016; Marder and Bucher 2007; Stein 2009).

Modulatory inputs from higher order ganglia modulate the pyloric and gastric mill networks to produce different rhythmic outputs (Nusbaum 2008; Stein 2009). The sensitivity of the *C. borealis* STG networks to many neuromodulators correlates with a varied diet of this species (Dickinson et al. 2008; Donahue et al. 2009; Stehlik 1993). Particularly relevant to this study, flexibility of STNS network output in multiple crab and lobster species includes neuronal switching between pyloric, gastric mill, and swallowing networks (Blitz et al. 2019; Fahoum and Blitz 2021; Hooper and Moulins 1990; Meyrand et al. 1994).

A modulatory projection neuron, modulatory commissural neuron 5 (MCN5), uses the neuropeptide Gly^1^-SIFamide to elicit a particular gastric mill motor pattern, including interactions between the pyloric and gastric mill networks that differ from those during activation of other modulatory inputs (Blitz et al. 2019; Fahoum and Blitz 2021). As part of its actions, MCN5 stimulation, or bath-applied Gly^1^-SIFamide, switches the lateral posterior gastric (LPG) neuron from solely being active with the pyloric rhythm, to dual-frequency bursting in time with the pyloric and gastric mill rhythms (Blitz et al. 2019; Fahoum and Blitz 2021). In *C. borealis*, LPG typically generates fast (∼1 Hz) bursts due to electrical coupling with the other pyloric pacemaker ensemble neurons (Blitz et al. 2019; Fahoum and Blitz 2021; Shruti et al. 2014). However, in the MCN5/Gly^1^-SIFamide modulatory state, LPG generates pyloric-timed bursts, due to continued electrical coupling to pyloric pacemaker neurons, plus slower, gastric mill-timed bursts that are generated due to Gly^1^-SIFamide modulation of LPG intrinsic properties (Blitz et al. 2019; Fahoum and Blitz 2021). The membrane properties underlying the switch from single to dual-network activity have not been determined (Fahoum and Blitz 2021).

Here, we extended our previous study to examine intrinsic properties modulated by Gly^1^-SIFamide, and ionic currents contributing to the LPG switching neuron generating intrinsic oscillations at a second frequency.

## Methods

### Animals

Male *C. borealis* crabs were purchased from the Fresh Lobster Company (Gloucester, MA) and housed in artificial seawater tanks (10-12°C). Immediately prior to dissection, crabs were anesthetized by packing in ice (45-55 min). Dissections were performed as previously described (Fahoum and Blitz 2021; Gutierrez and Grashow 2009). The foregut was removed from the crab, bisected ventrally, and pinned in a Sylgard (170: Thermo Fisher Scientific, Watham, MA)-coated dissection dish containing chilled (8- 10°C) *C. borealis* physiological saline. The STNS was dissected free from muscle and other tissue, transferred to a petri dish and pinned on a layer of clear Sylgard (184; Thermo Fisher Scientific). Preparations were stored overnight at 4°C in *C. borealis* physiological saline until used in electrophysiology experiments the following day.

### Solutions

*C. borealis* physiological saline (pH 7.4 – 7.6) consisted of the following (in mM): 440 NaCl, 26 MgCl_2_, 13 CaCl_2_, 11 KCl, 10 Trisma base, 5 Maleic acid. For low Ca^2+^ saline, 90% of Ca^2+^ was replaced with equimolar (11.7 mM) MnCl_2_. Riluzole (BD Solutions), a persistent sodium current blocker (Tohidi and Nadim 2009), was dissolved in 100% dimethylsulfoxide (DMSO; Thermo Fisher Scientific) at 100 mM and aliquots stored at - 20°C. Prior to use, riluzole aliquots were thawed and diluted in *C. borealis* saline (30 µM) such that the final DMSO concentration was less than 1%. At this concentration, DMSO does not affect STG neurons (Pérez-Acevedo and Krenz 2005; Scholz et al. 2001; Stein et al. 2005). Cesium chloride (CsCl; Sigma Aldrich), a hyperpolarization-activated inward current blocker (Peck et al. 2006; Tohidi and Nadim 2009; Zhu et al. 2016), was dissolved in *C. borealis* saline on the day of experiments at a final concentration of 5 mM. Picrotoxin (PTX; Sigma Aldrich) was dissolved in *C. borealis* saline or low Ca^2+^ saline at 10 µM and mixed thoroughly (> 45 min) immediately prior to use. PTX was used to block inhibitory glutamatergic synapses (Fahoum and Blitz 2021; Marder and Eisen 1984). The neuropeptide Gly^1^-SIFamide (GYRKPPFNG-SIFamide, custom peptide synthesis: Genscript) (Blitz et al. 2019) was dissolved in H_2_0 Optima (10^-2^ M) and stored at -20 °C. Aliquots were thawed and dissolved directly in the appropriate solution (*C. borealis* saline, 10 µM PTX in *C. borealis* saline or low Ca^2+^ saline, or saline plus CsCl and/or Riluzole) immediately prior to use at a final concentration of 5 µM. Solution changes were controlled by a switching manifold. Squid internal electrode solution consisted of (in mM): 20 NaCl, 15 Na_2_SO_4_, 10 Hepes, 400 potassium gluconate, 10 MgCl_2_ (Hooper et al. 2015).

### Electrophysiology

Petroleum jelly (Vaseline; CVS) wells were built around nerves to electrically isolate them from the rest of the bath. Extracellular recordings were obtained using paired stainless-steel electrodes. One electrode from each pair was placed adjacent to the nerve within a petroleum jelly well and the reference electrode from each pair was placed outside the well. Extracellular recordings were amplified with a differential AC amplifier (Model 1700; A-M systems, Sequim, WA).

The STG was desheathed prior to each experiment to facilitate visualizing STG somata for impalement. Intracellular recordings were obtained using pulled-glass sharp microelectrodes (30-50 MΩ) filled with a squid internal electrode solution (*see Solutions*). Intracellular recordings were amplified with an Axoclamp 900A amplifier (Molecular Devices, San Jose, CA). Intracellular and extracellular signals were digitized (∼5 kHz sampling rate) and recorded to a lab computer (Dell, Round Rock, TX) using data acquisition hardware (Micro1401; Cambridge Electronic Design [CED], Cambridge, U.K.) and software (Spike2; CED). Current injections were performed manually through the Axoclamp 900A commander software or via custom Spike2 stimulus sequence files. Intracellularly recorded neurons were identified by their activity patterns, synaptic input, and by confirming their axonal projection pattern through spike-triggered averaging of neuron and nerve recordings (Fahoum and Blitz 2021; Hudson et al. 2010; Marder and Bucher 2007). For accurate membrane potential measurements during current injections, some experiments were performed with either two-electrode current clamp (TECC; dual-impalement) or discontinuous current clamp (DCC; single impalement; sampling rate: 5 kHz). For DCC, the electrode time constant was minimized using the internal amplifier capacitance neutralization. In some experiments, electrodes were coated with Sylgard 184 (Fisher Scientific) to reduce electrode capacitance and facilitate DCC (Opdyke and Calabrese 1995).

Electrophysiology experiments, unless otherwise specified, were performed in the presence of PTX with both PD neurons hyperpolarized (PTX:PD_hype_) by negative current injection to eliminate the influence of rhythmic input through electrical synapses between LPG and the AB/PD pyloric pacemaker ensemble (Fahoum and Blitz 2021; Marder et al. 2017; Shruti et al. 2014). PTX eliminates the influence of glutamatergic chemical synapses, including the single known chemical synaptic input to LPG, the LP neuron (Fahoum and Blitz 2021; Marder and Eisen 1984). In a subset of experiments without PTX, hyperpolarizing DC current was injected into the LP neuron (LP_hype_) to eliminate input from LP to LPG (Fahoum and Blitz 2021).

### LPG Burst Detection

To identify LPG bursts, LPG spikes were detected in extracellular nerve (*lpgn*) recordings. For the PTX:PD_on_ condition, a histogram of LPG interspike intervals (ISIs) across Gly^1^-SIFamide applications in each experiment. Histograms included a cluster of short duration ISIs representing intervals between spikes within a burst and a cluster of longer duration ISIs representing the intervals between bursts. The median of the two ISI distribution peaks was calculated and selected as the maximum interval for two successive spikes to be considered part of the same burst. After calculating an ISI cutoff value, LPG bursts were identified as groups of two or more successive spikes with ISIs shorter than the calculated ISI cutoff (Fahoum and Blitz 2021). LPG slow bursts were then selected by subtracting LPG bursts with a duration shorter than two pyloric cycles (two consecutive PD bursts), using a custom Spike2 script (Fahoum and Blitz 2021).

This eliminated the faster, pyloric-timed LPG bursts originating from electrical coupling to the AD/PD neurons (Shruti et al. 2014). All LPG burst analysis in the PTX:PD_on_ condition includes only the slower, intrinsic LPG bursts. For the PTX:PD_hype_ condition a cutoff ISI was used to identify bursts as described for the PTX:PD_on_ condition. However, all LPG bursts during PTX:PD_hype_ were considered to be intrinsically generated and were included in analyses because LPG was isolated from rhythmic synaptic input from both networks.

### Analysis of Intrinsic Properties

Input resistance was calculated from the change in voltage in response to hyperpolarizing current injections (-1 nA; 5 s) from the same membrane potential in PTX:PD_hype_ and 5 µM Gly^1^-SIFamide in PTX:PD_hype_. Sag potential amplitude was measured as the difference between the membrane potential immediately before the end of a hyperpolarizing current injection and the most hyperpolarized membrane potential near the beginning of a hyperpolarizing current injection. In each experiment the current injection amplitude was adjusted in the SIF:PTX:PD_hype_ condition to match the most hyperpolarized membrane potential (± 4.0 mV) reached in the PTX:PD_hype_ condition in response to -1 nA (5 s). The amplitude measured during two current injections per condition, per experiment were averaged. To quantify post-inhibitory rebound, the four second period immediately preceding the same hyperpolarizing current steps used to measure sag was selected as a baseline. This baseline was subtracted from the four second period beginning 200 ms after the termination of the current step, and the integrated area of the LPG membrane potential above baseline was calculated using the Spike2 ‘Area Under Curve’ function (Angstadt et al. 2005; Blitz 2017). Two measurements in the PTX:PD_hype_ condition and in the SIF:PTX:PD_hype_ condition were averaged.

Firing frequency (# of spikes – 1 / burst duration) was measured across 5 s depolarizing current steps (0.25-3.0 nA). A subset of these current steps was used to quantify spike frequency adaptation (SFA). In each experiment, the smallest current amplitude that elicited spiking across the 5 s duration in the PTX:PD_hype_ condition (1 – 2.5 nA), and the current amplitude that depolarized LPG to the same membrane potential (± 3.0 mV) in the SIF:PTX:PD_hype_ condition (1-2.5 nA) were selected. The ISIs across each step were plotted as a function of time and a linear line of best fit was calculated in Excel (Microsoft, Redmond, WA). The slope value of the best fit line was used as a measure of SFA as it represented the rate at which the interval between spikes changed across the current step.

### Analysis of Burst Properties

To quantify the influence of ion channel blockers and ion substitution, LPG bursts were analyzed from extracellular recordings. Burst duration (interval from first to last spike within a burst; s), cycle period (interval from first spike in a burst to the first spike in the subsequent burst; s), and interburst interval (IBI; cycle period – burst duration; s) was measured for LPG slow bursts across a 200 s window once Gly^1^-SIFamide effects reached a steady-state, (stable effects for at least 200 s), beginning immediately after Gly^1^-SIFamide effects reached steady-state. Trough voltage was measured from intracellular recordings as the most hyperpolarized membrane potential (mV).

### Statistical Analysis

Following quantification, data was evaluated for normality using a Shapiro-Wilk normality test to determine whether a parametric or non-parametric test should be used. For comparisons between two groups, a paired t-test or Wilcoxon signed rank test was used. This analysis was performed to compare sag potential, input resistance, integrated area under the curve, and ISI rate of change between PTX:PD_hype_ and SIF:PTX:PD_hype_ conditions. For each category of experiments, a statistical power analysis was performed and a power level ≥ 0.8 was considered sufficient. For some data sets, power was below 0.8 (Table 1). In these datasets there was high variability at n values (n = 5 – 8) comparable to other data sets in this study (e.g., Fig. 5, 6) which would require extremely large n values to generate a sufficient power level. Based on visual observation of plots of all individual experiments in these data sets and the large n values that would be required, we decided it was not prudent to pursue these experiments further. A two-factor within-subjects repeated measures (RM) ANOVA was used to measure the effects of current amplitude and Gly^1^-SIFamide on LPG firing frequency, followed by Tukey pairwise comparisons at each current level in PTX:PD_hype_ SIF:PTX:PD_hype_. For pharmacological blockade studies, LPG activity parameters were compared across three conditions: pre-control (SIF:PTX:PD_hype_), treatment (SIF:PTX:PD_hype_ + blocker), and post-control (SIF:PTX:PD_hype_). To compare the three conditions, a one-way RM ANOVA was used followed by post-hoc pairwise comparisons where appropriate (Bonferroni t-test). Statistical significance was p < 0.05. Statistical analyses were performed using SigmaPlot software (v. 14; Systat Software, San Jose, CA) or SAS (v. 9.4; Cary, NC). Excel software was used to organize data and MATLAB (Mathworks, Natick, MA) and CorelDraw (Corel Corporation, Ontario, CA) were used to generate graphs and figures. Scripts/code available at https://github.com/blitzdm/Snyder_Blitz_2022.

**Table 1.** LPG intrinsic bursts were altered by ion channel blockers.

| Blocker | PDs | Parameter | Mean ± SEM | | | $F^a$ | Df<sup>b</sup> | $p$-value | | | Power |
| --- | --- | --- | --- | --- | --- | --- | --- | --- | --- | --- | --- |
|  |  |  | Pre | Blocker | Post |  |  | Pre vs Blocker | Post vs Blocker | Pre vs Post |  |
| CsCl<br>(n = 6) | Hype | Cycle Pd<sup>c</sup> | 12.6 ± 1.99 | 25.78 ± 4.17 | 12.04 ± 2.77 | 7.479 | 5,2 | 0.022 | 0.013 | 0.590 | 0.809 |
| | | Burst Dur. | 6.24 ± 0.97 | 5.51 ± 1.10 | 5.48 ± 1.81 | 0.143 | 5,2 | *n.s.* | *n.s.* | *n.s.* | 0.050 |
| | | IBI | 6.42 ± 1.33 | 20.28 ± 4.04 | 5.55 ± 1.60 | 20.688 | 5,2 | <0.001 | <0.001 | 0.743 | 0.999 |
| CsCl<br>(n = 8-9) | On | Cycle Pd | 13.49 ± 1.59 | 21.13 ± 1.94 | 12.28 ± 2.35 | 6.757 | 7,2 | 0.022 | 0.013 | 0.649 | 0.781 |
| | | Burst Dur. | 5.30 ± 0.64 | 3.72 ± 0.26 | 4.94 ± 1.59 | 0.809 | 8,2 | *n.s.* | *n.s.* | *n.s.* | 0.050 |
| | | IBI | 7.85 ± 1.21 | 17.44 ± 2.17 | 7.34 ± 1.16 | 13.386 | 7,2 | 0.001 | 0.001 | 0.820 | 0.984 |
| Riluzole<br>(n = 6) | Hype | Cycle Pd | 12.30 ± 1.43 | 6.50 ± 0.44 | 7.72 ± 0.87 | 12.268 | 5,2 | 0.004 | 0.463 | 0.022 | 0.943 |
| | | Burst Dur. | 5.75 ± 0.54 | 1.65 ± 0.31 | 2.61 ± 0.16 | 23.698 | 5,2 | <0.001 | 0.234 | 0.005 | 0.999 |
| | | IBI | 6.55 ± 1.10 | 4.85 ± 0.36 | 5.11 ± 0.72 | 1.834 | 5,2 | *n.s.* | *n.s.* | *n.s.* | 0.147 |

<sup>a</sup>One-Way RM ANOVA, followed by Holm-Sidak post hoc pairwise comparisons. <sup>n.s.</sup>Post hoc was not performed when $p > 0.05$ . <sup>b</sup>Degrees of freedom (Df) reported for between subjects, within subjects. <sup>c</sup>This data set was transformed by log10 to normalize data prior to statistical testing.

## Results

### Modulator-activated Intrinsic LPG Bursting

To examine intrinsic properties and mechanisms of burst generation, we isolated LPG neurons from rhythmic input from the pyloric and gastric mill networks. Picrotoxin (PTX, 10 µM) blocks glutamatergic inhibition (Bidaut 1980; Cleland and Selverston 1995; Marder and Eisen 1984), including the only chemical synapse onto LPG from other pyloric neurons. In Saline:PTX, LPG continues to produce pyloric-timed oscillations due to electrical coupling with the AB/PD neurons (Fig. 1B-C) (Bartos et al. 1999; Marder et al. 2017). When Gly^1^-SIFamide (5 µM) is applied in the presence of PTX (SIF:PTX), LPG generates dual-network activity including faster, shorter duration bursts in time with the pyloric rhythm (Fig. 1D, red boxes), and slower, longer duration gastric-mill timed bursts (Fig. 1D, blue box). Although the gastric mill-timed LPG bursting occurs via intrinsic mechanisms (Fahoum and Blitz 2021), LPG may receive synaptic input from other gastric mill neurons in order to coordinate their activity (S-RH Fahoum, DM Blitz, unpublished). Any such influence was removed in these experiments via PTX, as PTX also blocks potential chemical synaptic input from other gastric mill neurons (Fahoum and Blitz 2021; Marder and Bucher 2007). To block the influence of rhythmic pyloric input, the two PD neurons were hyperpolarized sufficiently to eliminate rhythmic bursting in the pyloric pacemaker (AB) neuron (Bartos et al. 1999; Blitz et al. 2008; Fahoum and Blitz 2021) (see Methods). Hyperpolarizing the PD neurons does not eliminate LPG slow intrinsic bursting due to the rectifying nature of the electrical coupling between PD and LPG (Fig. 1B) (Fahoum and Blitz 2021; Shruti et al. 2014).

**Figure 1:**
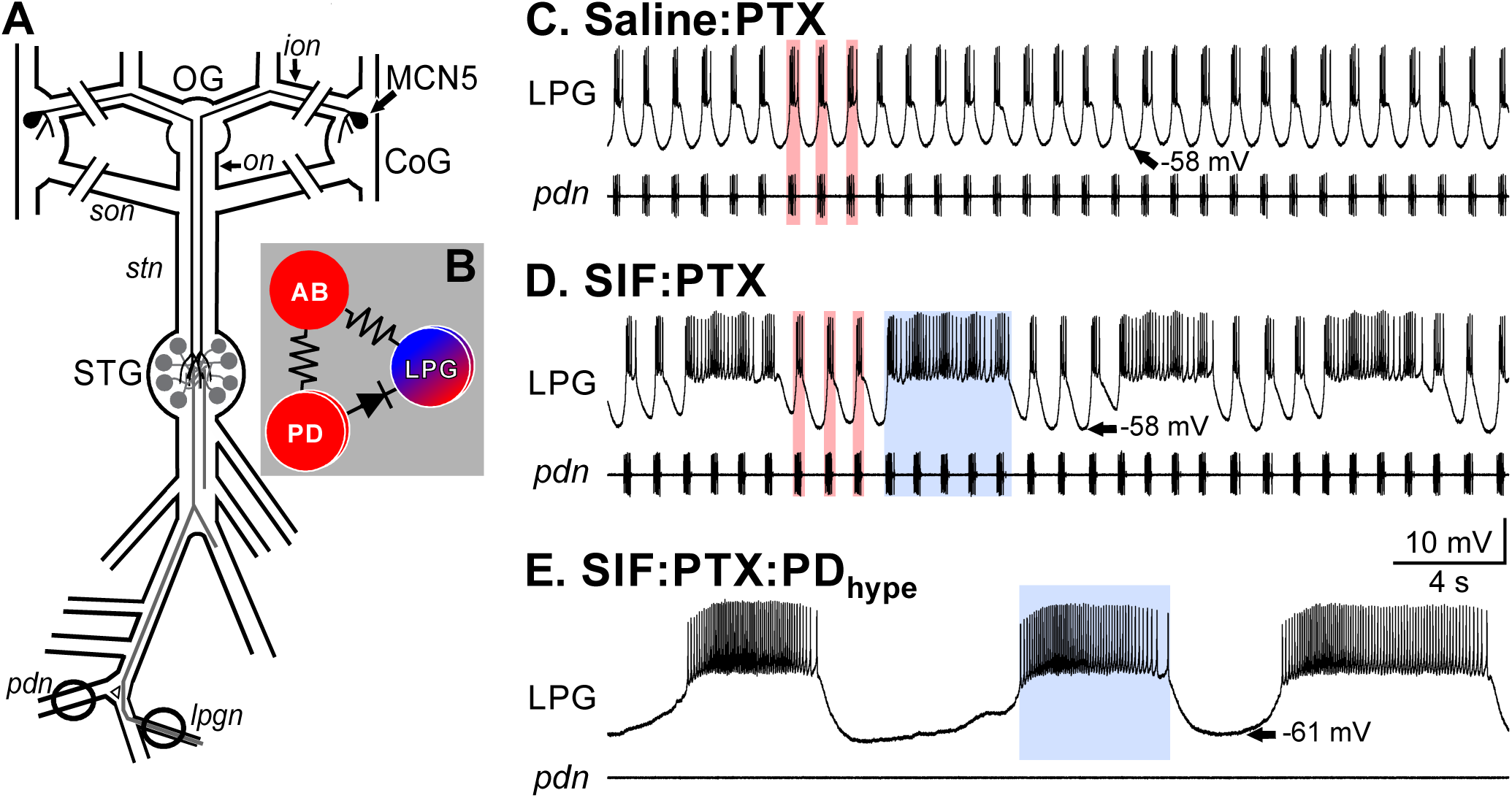
Modulatory neuron MCN5 or bath application of its neuropeptide Gly^1^- SIFamide elicits a switch in LPG neuron activity from only pyloric-timed oscillations to dual pyloric/gastric mill-timed oscillations. A) The isolated stomatogastric nervous system consists of the paired CoGs and single OG from which modulatory neurons originate, and the single STG containing pyloric and gastric mill network neurons. This study used intracellular recordings from network neurons in the STG and extracellular recordings from peripheral nerves (circles on nerves). There is a single MCN5 soma in each CoG which project an axon through the *ion* and *stn* that terminates in the STG. Double lines through *ions* and *sons* indicate cuts made to eliminate modulatory input to the STG from MCN5 and other modulatory neurons located in the CoGs. B) The rhythmogenic pyloric pacemaker ensemble located within the STG consists of a single AB neuron, two PD neurons and two LPG neurons coupled through electrical synapses (resistors, electrical synapses; diode, rectifying electrical synapse). Red circles indicate neurons solely active in the pyloric rhythm, red/blue circles indicate that LPG can participate in both the pyloric and gastric mill networks. C) Baseline conditions for this study included the presence of the inhibitory glutamate channel antagonist picrotoxin (PTX, 10 µM) to block chemical synaptic input to LPG. In this condition, AB (not shown), PD (*bottom*, extracellular *pdn* recording), and LPG (*top*, intracellular recording) generate rhythmic pyloric-timed (∼1 Hz) oscillations (red boxes) due to the intrinsic properties of the AB neuron and the electrical coupling among this pacemaker ensemble (Marder and Eisen 1984; Shruti et al. 2014). In this condition, the gastric-mill rhythm is inactive (not shown). D) In 5 µM Gly^1^-SIFamide in PTX, LPG exhibits dual frequency bursting in time with the fast pyloric (∼1 Hz; red boxes) and slow gastric mill (∼0.1 Hz; blue box) rhythms. E) LPG slow bursts (blue box) occur via intrinsic properties and therefore persist in 5 µM Gly^1^-SIFamide in PTX with hyperpolarizing current in the PD neurons (PTX:PD_hype_) to silence the pyloric rhythm (Fahoum and Blitz 2021). Scale bars apply to C-E. All recordings are from the same preparation. Abbreviations: Ganglia: CoG, commissural ganglion; OG, Oesophageal ganglion; STG, Stomatogastric ganglion. Nerves: *ion*, inferior oesophageal nerve; *lpgn*, lateral posterior gastric nerve; *on*, oesophageal nerve; *pdn*, pyloric dilator nerve; *son*, superior oesophageal nerve; *stn*, stomatogastric nerve. Neurons: AB, anterior burster; LPG, lateral posterior gastric; MCN5, modulatory commissural neuron 5; PD, pyloric dilator.

Thus, in this isolated condition without rhythmic input from either circuit, Gly^1^-SIFamide (SIF:PTX:PD_hype_) elicits intrinsic bursting at a slow, gastric mill frequency (∼ 0.06 Hz) (Fig 1E) (Fahoum and Blitz 2021). The two baseline conditions used throughout this study were PTX:PD_on_ to examine dual-frequency bursting (e.g., Fig. 1D) and PTX:PD_hype_ to examine isolated slow bursting (e.g., Fig. 1E).

### Intrinsic Excitability

As an initial measure of LPG excitability, we examined LPG input resistance (R_i_) near its resting potential. Injecting negative current (-1 nA; 5 s) in the PTX:PD_hype_ condition hyperpolarized LPG for the duration of the current step (Fig. 2A; grey trace). In the presence of Gly^1^-SIFamide (5 µM), an equivalent current step from the same initial membrane potential brought LPG to a more hyperpolarized potential (Fig. 2A; black trace), indicating an increased input resistance in Gly^1^-SIFamide (5 µM; SIF:PTX:PD_hype_) compared to PTX:PD_hype_ alone (Fig. 2A-B; initial V_m_: PTX:PD_hype_, -48.9 ± 2.0 mV; SIF:PTX:PD_hype_, -48.7 ± 1.9 mV; R_i_: PTX:PD_hype_, 19.2 ± 2.1 MΩ; SIF:PTX:PD_hype_, 22.6 ± 2.4 MΩ; n = 11; *t*_(10)_ = 4.462, p = 0.0012, paired *t*-test). Thus, Gly^1^-SIFamide modulation increased the input resistance of LPG near – 50 mV, indicating increased excitability within the voltage range of its typical pyloric oscillations.

**Figure 2:**
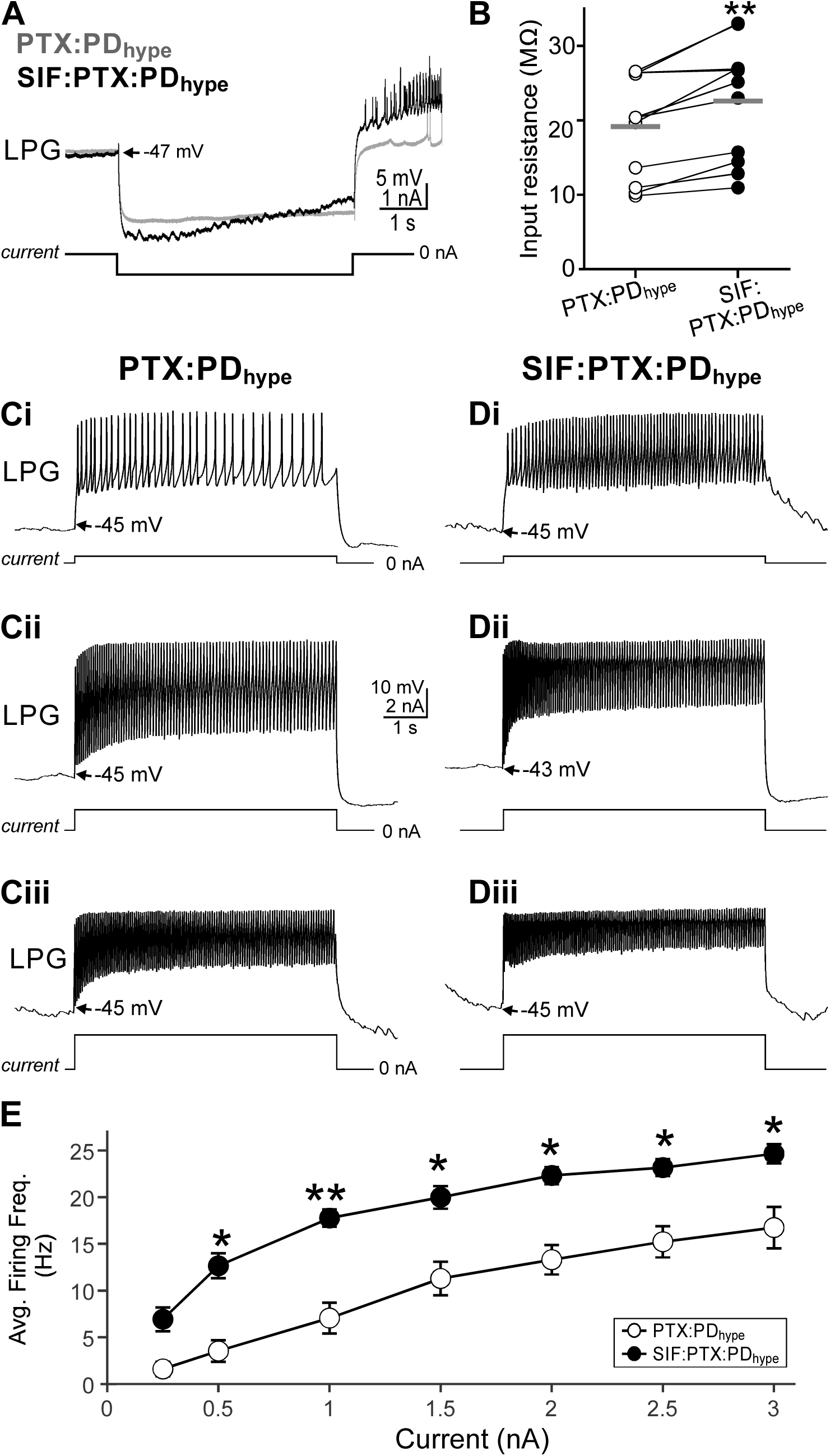
LPG excitability was increased by Gly^1^-SIFamide. A) LPG input resistance was measured with a 1 nA hyperpolarizing current injection starting near LPG resting potential in the baseline PTX:PD_hype_ condition and matched to the same initial membrane potential in the SIF:PTX:PD_hype_ condition. In an example experiment, LPG was hyperpolarized more in Gly^1^-SIFamide (5 µM; SIF:PTX:PD_hype_; black trace) compared to PTX:PD_hype_ (grey trace). Each trace is the average of three trials. B) Plot of input resistance in PTX:PD_hype_ (open circles) and SIF:PTX:PD_hype_ (filled circles). Each line with pair of symbols is a single experiment. Gray bars indicate averages for each condition. (** p<0.01; n = 11; paired t-test). C-D) LPG was depolarized with a 0.5, 1.5, and 2.5 nA current injection for 5 s in PTX:PD_hype_ (Ci-iii) or SIF:PTX:PD_hype_ (Di-Diii). The pre-current step membrane potential was matched between the two conditions using current injection in DCC mode (see Methods). Scale bars between Cii and Dii apply to all traces in C-D. All traces in C-D are from the same experiment. E) The average (± SEM) LPG firing frequency across each level of current injection is plotted for PTX:PD_hype_ (open circles) and for SIF:PTX:PD_hype_ (filled circles). (0.5 – 3.0 nA; *p<0.05, **p<0.01; n = 4-10; Two-way RM ANOVA, Tukey post-hoc).

To further examine excitability across a larger range of voltages, the LPG firing frequency was measured before and during Gly^1^-SIFamide application in response to depolarizing current steps (0.25 to 3 nA; 5 s) from the same membrane potential (± 3.0 mV). In an example experiment, in the PTX:PD_hype_ condition, a depolarizing current step (0.5 nA; 5 s) evoked a relatively low firing frequency (7 Hz) (Fig. 2Ci), while 1.5 and 2.5 nA elicited higher firing rates (18 and 22 Hz) (Fig. 2Cii, Ciii). In the presence of Gly^1^-SIFamide, the same amplitude current injections elicited higher firing frequencies (15, 25, 26 Hz; Fig. 2Di-Diii). Across experiments, the frequency/current (F/I) relationship was altered by Gly^1^-SIFamide, as a function of current step amplitude (Fig. 2E) (Interaction effects: Current step amplitude and PTX versus Gly^1^-SIFamide; *F*_(6, 42)_ = 2.999, p = 0.016, Two Way RM ANOVA). Specifically, although there was no difference in firing frequency at 0.25 nA (*t*_(6.99)_ = -1.83; p = 0.109, Tukey post-hoc comparison) the firing frequency was higher in Gly^1^-SIFamide compared to PTX:PD_hype_ in response to current steps from 0.5 to 3 nA (**0.5 nA**, *t*_(6.41)_ = -3.49, p = 0.012; **1.0 nA**, *t*_(6.41)_ = -4.17, p=0.005; **1.5 nA,** *t*_(6.57)_ = -3.46, p = 0.012; **2.0 nA**, *t*_(6.41)_ = -3.45, p = 0.012; **2.5 nA**, *t*_(6.75)_ = -2.89, p = 0.024; **3.0 nA**: *t_(_*_8.66)_ = -2.66, p = 0.027; n = 4 – 10), The positive shift in the LPG F/I relationship indicates that LPG excitability was increased by Gly^1^-SIFamide within its typical membrane potential range during an ongoing pyloric rhythm.

### Intrinsic Membrane Properties

During the current injections used to assess the LPG F/I relationship, we noted that firing frequency appeared to wane across 5 s current steps (e.g., Fig. 2Ci). Thus, we examined the property of spike frequency adaptation (SFA), in which the interspike interval progressively increases in duration across a period of activity (Ha and Cheong 2017; Katz and Frost 1997; Madison and Nicoll 1982). To test SFA in LPG, we used current steps (1 – 2.5 nA, 5 s) that depolarized LPG to the same membrane potential before and during Gly^1^-SIFamide application (PTX:PD_hype_, -31.7 ± 2.3 mV; SIF:PTX:PD_hype_, -29.5 ± 2.0 mV). In the PTX:PD_hype_ condition a depolarizing current step (1 nA; 5 s) elicited an initial LPG firing frequency of ∼ 22 Hz (initial 0.5 s; Fig.3A, Ai) which decayed to ∼ 8 Hz (final 0.5 s; Fig. 3A, Aii). When LPG was depolarized to the same membrane potential in Gly^1^-SIFamide, the initial LPG firing frequency (initial 0.5 s,

**Figure 3:**
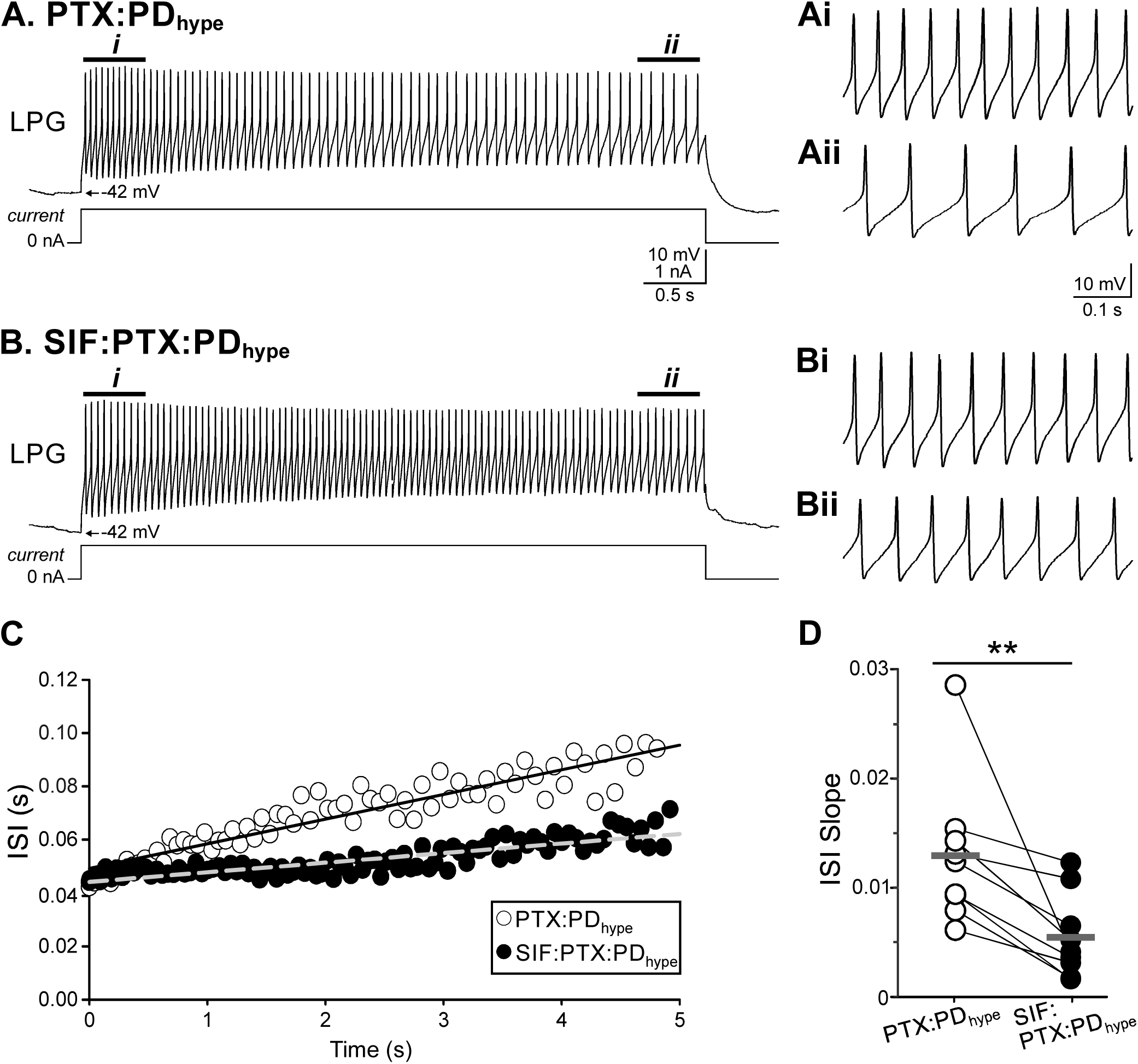
LPG spike frequency adaptation was decreased by Gly^1^-SIFamide. LPG firing frequency was assessed across a 5 s (1.0 nA) current injection in the PTX:PD_hype_ (A) and SIF:PTX:PD_hype_ (B) conditions. (Ai-ii; Bi-ii) Expanded sections from A, B highlight the firing frequency at the beginning (*i*; 0.5 s) and end (*ii*; 0.5 s) of the current step. Scale bar in A applies to A and B. Scale bar in Ai applies to Ai-Bii. All recordings are from the same preparation. C) Plot of interspike interval versus time for all intervals in the current steps from A (open circles; PTX:PD_hype_) and B (filled circles; SIF:PTX:PD_hype_). Linear fits to the data are indicated by the solid (PTX:PD_hype_) and grey dashed (SIF:PTX:PD_hype_) lines. (D) The slope of each line across experiments was then plotted for PTX:PD_hype_ (open circles) and SIF:PTX:PD_hype_ (filled circles) . Each line plus set of symbols plots the ISI slopes for a single experiment. Gray bars indicate averages for each condition. (**p = 0.004; n = 9; Wilcoxon signed-rank test).

∼24 Hz; Fig. 3B, Bi) was similar to PTX:PD_hype_. However, the firing frequency did not decay to the same extent (final 0.5 s, ∼14 Hz; Fig. 3B, Bii) as it did in the control PTX:PD_hype_ condition. To quantify the change in firing frequency, we plotted the interspike interval (ISI) as a function of time in PTX:PD_hype_ (open circles) and PTX:PD_hype_ plus Gly^1^-SIFamide (filled circles) (Fig. 3C) and fit each data set with a line of best fit. In this example, the slope of the line of best fit was steeper in PTX:PD_hype_, indicating a greater amount of SFA in this control condition compared to Gly^1^-SIFamide application (Fig. 3C; Slope: PTX:PD_hype_, 0.0093; SIF:PTX:PD_hype_, 0.0036). ISIs were plotted in this manner for each experiment and the slopes of the fit lines were used to compare SFA between conditions. As plotted in figure 3D, the ISI/time slope was steeper in PTX:PD_hype_ versus SIF:PTX:PD_hype_ (Slope PTX:PD_hype_, 0.013 ± 0.002; SIF:PTX:PD_hype_, 0.005 ± 0.001; n = 9; Z = -2.666, p = 0.004, Wilcoxon signed-rank) (Fig. 3D). Thus Gly^1^-SIFamide decreased the extent of SFA in LPG, which can delay burst termination (Bucher et al. 2015; Ha and Cheong 2017).

As plateau properties can contribute to the active phase of bursting (Bucher et al. 2015; Tong and McDearmid 2012), we asked whether LPG expresses plateau potentials. We tested for plateau potentials using brief (10 – 200 ms) depolarizing (1 – 10 nA) current injections. Starting at membrane potentials between -45 and -60 mV, these brief depolarizing steps did not elicit plateau potentials in control PTX:PD_hype_ conditions or during Gly^1^-SIFamide (5 µM in PTX:PD_hype_) application (n = 6; data not shown). Additionally, if plateau properties contributed to slow bursts, brief hyperpolarizing current injections of sufficient amplitude should prematurely terminate these bursts. However, hyperpolarizing current injections (10 – 200 ms) across a range of amplitude (-1 to -10 nA), did not terminate Gly^1^-SIFamide-elicited bursting in PTX:PD_hype_ (n = 6; data not shown).

Another intrinsic membrane property that often contributes to oscillatory activity in many neuron types is post-inhibitory rebound (PIR), in which a period of inhibition triggers a more depolarized membrane potential, often with increased activity, compared to pre-inhibition (Angstadt et al. 2005; Blitz 2017; Bucher et al. 2015; Picton et al. 2018). In control conditions (PTX:PD_hype_), following a negative current step (-1 nA; 5 s; trough V_m_: -66 mV) the LPG membrane potential depolarized above the baseline potential (grey dashed line), and fired several (12) action potentials before returning to baseline in the example shown (Fig. 4Ai). This rebound response indicates that LPG expressed PIR in the control condition. After being hyperpolarized to a comparable membrane potential (trough V_m_: -67 mV; -1 nA; 5 s) in Gly^1^-SIFamide, the LPG membrane potential depolarized to a greater extent and LPG fired more action potentials (61; Fig. 4Aii) compared to PTX:PD_hype_, suggesting that PIR is modulated by Gly^1^-SIFamide. To quantify PIR, we measured the integrated area under the LPG membrane potential during a 4 s window after the end of the current step (Fig. 4Ai-Aii; shaded area; see Methods). The integrated area was greater in Gly^1^-SIFamide (20.5 ± 3.0 mV*ms) compared to PTX:PD_hype_ (4.1 ± 1.0 mV*ms; n = 8; paired *t*-test, *t*_(7)_ = -6.356, p = 0.00038) (Fig. 4B). Additionally, more action potentials were elicited in the same 4 s window post-hyperpolarization in SIF:PTX:PD_hype_ compared to the PTX:PD_hype_ control (PTX:PD_hype_: 2.44 ± 1.44 spikes; SIF:PTX:PD_hype_ 26.38 ± 6.77 spikes; n = 8; paired *t*- test*, t*_(7)_ = -4.115, p = 0.004) (Fig. 4C).

**Figure 4:**
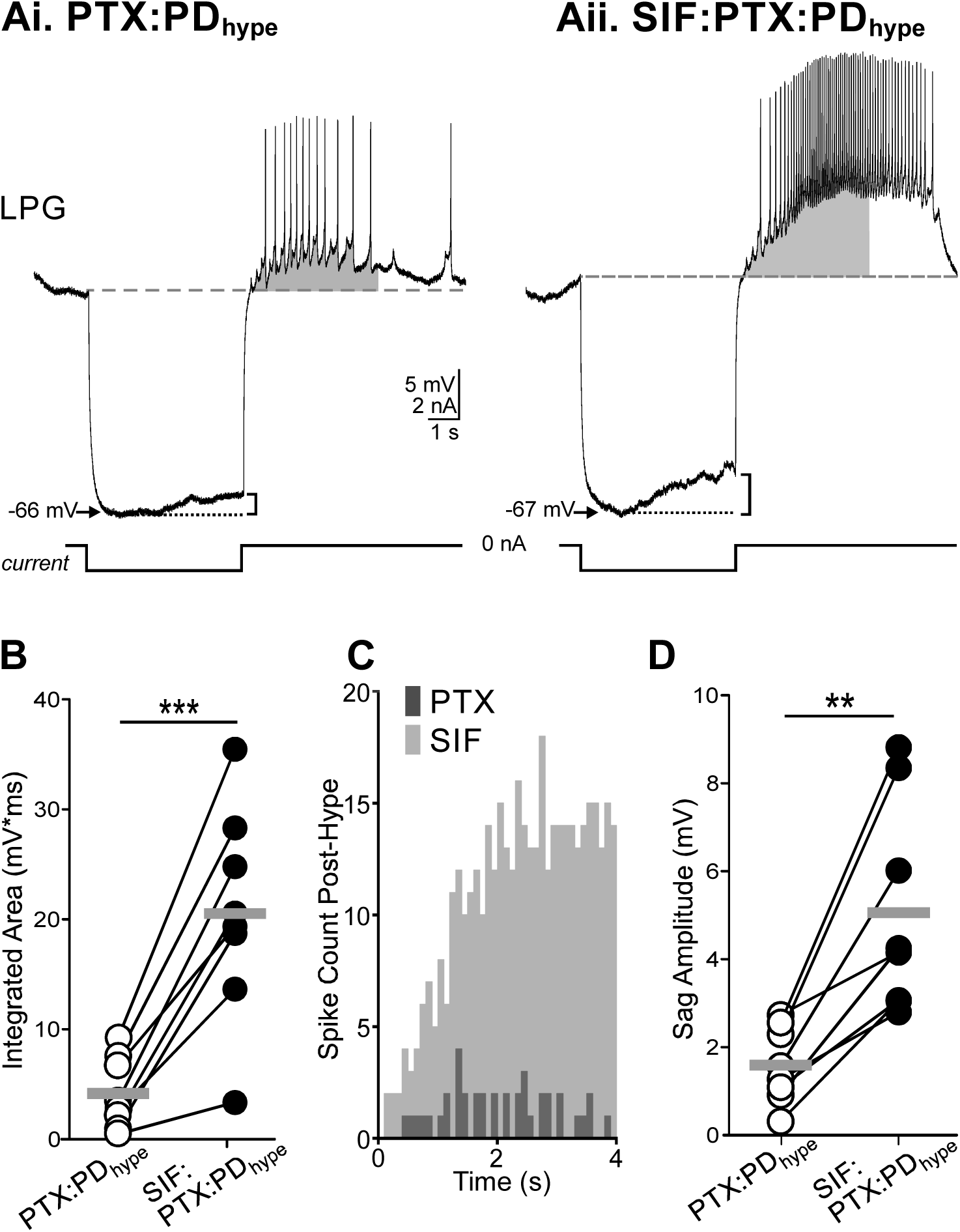
LPG post-inhibitory rebound was enhanced by Gly^1^-SIFamide. A hyperpolarizing current step was used to bring LPG to the same membrane potential in PTX:PD_hype_ (Ai) and SIF:PTX:PD_hype_ (Aii) for 5 s. The LPG membrane potential sagged upwards (bracket) toward the end of the current step. After the end of the current step, LPG generated a rebound above baseline (grey dashed line) including firing action potentials. The grey shaded areas indicate the analysis region for (B) (see Methods). Scale bars apply to Ai, Aii. Recordings are from the same preparation. B) Plot of integrated area under the curve for each experiment in the presence (filled circles) and absence (open circles) of Gly^1^-SIFamide (***p<0.001; n = 8; paired t-test). C) The cumulative number of spikes per bin is plotted across the 4 s window (grey areas in Ai, Aii) for PTX:PD_hype_ (dark grey) and SIF:PTX:PD_hype_ (lighter grey) (Bin size: 0.1 s). D) Plot of sag amplitude (change in membrane potential, see brackets in Ai, Aiil; mV) for each experiment in the presence (filled circles) and absence (open circles) of Gly^1^- SIFamide. (**p<0.01; n = 8; paired t-test). Gray bars in B, D indicate averages for each condition.

There are multiple ionic currents that can contribute to PIR, including the hyperpolarization-activated inward current, I_h_ (Bucher et al. 2015; Combe and Gasparini 2021; Peck et al. 2006; Picton et al. 2018). A depolarizing sag in membrane potential during maintained hyperpolarization suggests the activation of I_h_. To assess whether I_h_ might contribute to PIR, including the larger PIR in Gly^1^-SIFamide, we measured the change in membrane potential from the most hyperpolarized level (dotted lines; Fig. 4Ai-Aii) to the voltage at the end of the step (“sag”: brackets, Fig. 4Ai-Aii) during 5 s current steps. LPG was hyperpolarized to a comparable membrane potential (± 4 mV; see Methods) in PTX:PD_hype_ and SIF:PTX:PD_hype_ in each preparation. The amplitude of the hyperpolarization-activated sag was larger in Gly^1^-SIFamide (Fig. 4D; PTX:PD_hype_: 1.58 ± 0.31; SIF:PTX:PD_hype_: 5.06 ± 0.85 mV; paired *t*-test, n = 8; *t*_(7)_ = -5.074, p = 0.00144). These data implicate a hyperpolarization-activated inward current as a contributor to PIR in LPG and a target for Gly^1^-SIFamide modulation. To further examine contributions of ionic currents to the intrinsic, slow LPG bursting, we used blockers and ion-substitution in both the isolated (PTX:PD_hype_), and dual-network (PTX:PD_on_) conditions.

### Ionic Current Contributions

The I_h_ current is a common target of neuromodulation (Combe and Gasparini 2021; Harris-Warrick 2002; Zhang et al. 2003). As I_h_ was implicated in PIR in LPG due to the enhanced sag potential during hyperpolarizing pulses, we assessed a role of I_h_ in LPG intrinsic bursting by blocking this current using extracellular cesium (5 mM CsCl) (Peck et al. 2006; Tohidi and Nadim 2009; Zhu et al. 2016) (Fig. 5). When Gly^1^-SIFamide was bath-applied in CsCl:PTX:PD_hype_, LPG generated intrinsic bursts, suggesting that the I_h_ current is not necessary for intrinsic bursting (Fig. 5Aii). However, there were differences in Gly^1^-SIFamide-elicited bursts in CsCl compared to the control condition (SIF:PTX:PD_hype_). Although four bursts were generated during Gly^1^-SIFamide in the absence (Fig. 5Ai) or presence (Fig. 5Aii) of CsCl, the cycle period was longer in CsCl in the example shown, with a notable change in the interburst interval (Fig. 5Ai-Aii, black bars denote duration of first interburst interval in PTX:PD_hype_). Plotting the average cycle period (Fig. 5B), interburst interval (Fig. 5C) and burst duration (Fig. 5D) with the same y-axis scaling, an increased cycle period in CsCl appeared to be due entirely to an increased interval between bursts. CsCl altered LPG slow burst cycle period (One way RM ANOVA, F_(5,2)_ = 7.749, p = 0.009, n = 6) and interburst interval (One way RM ANOVA, F_(5,2)_ = 20.688, p < 0.001, n = 6), but did not change burst duration (One way RM ANOVA, F_(5,2)_ = 0.143, p = 0.868, n = 6). Specifically, Gly^1^-SIFamide-elicited intrinsic bursting cycle period was longer in the presence of CsCl compared to the SIF- Pre application (Fig. 5B; n = 6; p < 0.05, Table 1). The interburst interval was also longer in SIF-CsCl compared to SIF-Pre and SIF-Post (Fig. 5C; n = 6; p < 0.001, Table 1). Together, these results implicate a role for the I_h_ current in regulating the timing of LPG intrinsic bursts. Specifically, the variable effects on burst duration but consistent effects on interburst interval and cycle period indicate that I_h_ primarily contributes to initiating a subsequent burst, but it is not necessary for intrinsic bursting.

**Figure 5:**
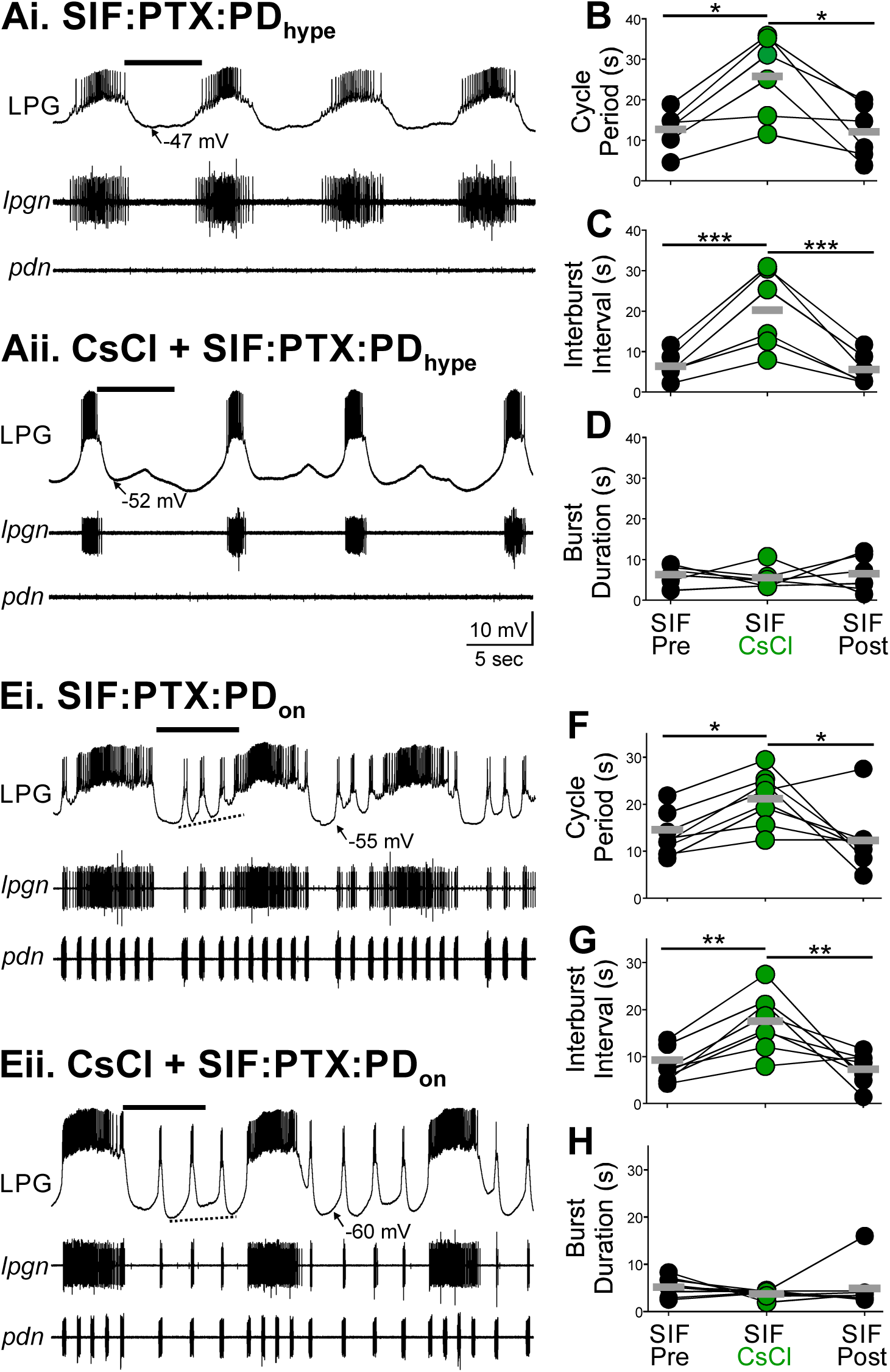
Blocking I_h_ current with CsCl increased the interburst interval in Gly^1^- SIFamide-elicited LPG intrinsic bursting. Ai) The LPG neurons (single LPG intracellular: top trace; paired LPGs, extracellular *lpgn:* middle trace) generated intrinsic bursts with the PD neurons hyperpolarized (extracellular *pdn:* bottom trace) during Gly^1^- SIFamide application (SIF:PTX:PD_hype_). (Aii) In the same preparation, LPG also generated slow bursting when Gly^1^-SIFamide was bath-applied in CsCl to block I_h_. Scale bars apply to Ai-Aii. The LPG slow burst cycle period (B), interburst interval (C) and burst duration (D) are plotted for SIF:PTX:PD_hype_ before (SIF Pre) and after (SIF Post) SIF:PTX:PD_hype_ in CsCl (SIF CsCl). Ei) In an example preparation, bath-applied Gly^1^-SIFamide in PTX with the pyloric rhythm intact (PTX:PD_on_) elicited periodic slow bursts that escape the pyloric rhythm and persisted for multiple pyloric cycles (PD neuron bursts, *pdn*). The membrane potential of pyloric-timed troughs in LPG between slow bursts progressively depolarized preceding a slow burst (dotted line). Eii) In the same preparation, application of Gly^1^-SIFamide with CsCl in the PTX:PD_on_ condition also elicited periodic slow bursts that escaped from the pyloric rhythm. In this condition, there was not a progressive depolarization of the trough membrane potential leading to the next slow burst (dotted line). Scale bars apply to Ei-Eii. Cycle period (F), interburst interval (G), and burst duration (H) during SIF:PTX:PD_hype_ before (SIF Pre), during (SIF CsCl), and after (SIF Post) SIF:PTX:PD_on_ in CsCl. Black bars above traces highlight the first interburst interval duration in SIF:PTX:PD_hype_ for comparison with the CsCl condition in A and E. In all graphs, each set of circles and lines connect data points from a single experiment and thick gray lines represent mean values for each condition. (SIF:PTX:PD_on_, black circles; CsCl + SIF:PTX:PD_on_, green circles). (B-D, n = 8; F-H, n = 7-8; *p<0.05, **p<0.01. ***p<0.001; one-way RM ANOVA, Holm-Sidak, see Table 1).

In control conditions, LPG must escape from the fast (∼1 Hz) pyloric-timed input to generate slow, intrinsic bursts. Generating a slow burst despite this rhythmic electrical coupling may have a different dependence on ionic currents compared to the isolated bursting examined above. We therefore tested the role of I_h_ in LPG slow bursting when it was receiving pyloric input (PTX:PD_on_). With an ongoing pyloric rhythm (*pdn*), LPG generated dual-frequency bursting in Gly^1^-SIFamide in a control SIF:PTX:PD_on_ condition (Fig. 5Ei), as well as in the presence of CsCl (5 µM) (Fig. 5Eii). Blocking I_h_ with the pyloric rhythm active also extended the Gly^1^-SIFamide-elicited LPG slow burst cycle period (Fig. 5E). Similar to the isolated LPG bursting, LPG slow bursts in the dual-frequency condition appeared to have longer cycle periods (Fig. 5F) that were primarily due to extended interburst intervals (Fig. 5E,G) and not due to a change in burst duration (Fig. 5H), evident in plots of all individual experiments with the same y-scaling for the three burst parameters (Fig. 5F-H). In the dual-frequency condition, CsCl changed LPG slow burst cycle period (One way RM ANOVA, F_(7,2)_ = 6.757, p = 0.009, n = 8), interburst interval (One way RM ANOVA, F_(7,2)_ = 13.386, p < 0.001, n = 8), but again did not alter burst duration (One way RM ANOVA, F_(7,2)_ = 0.809, p = 0.464, n = 8). Specifically, the cycle period was longer in CsCl + SIF:PTX:PD_on_ compared to SIF-Pre and SIF-Post (Fig. 5F; n = 8; p < 0.05, Table 1) and the interburst interval was longer in CsCl + SIF:PTX:PD_on_ compared to SIF:PTX:PD_on_ before and after the CsCl application (SIF-Pre and SIF-Post; Fig. 5G; n = 8; p < 0.001, Table 1). Thus, in both isolated LPG slow bursting and dual-frequency bursting, blocking I_h_ extended the interburst interval without a consistent effect on burst duration, but did not prevent slow bursting. These data suggest that I_h_, and likely PIR, contribute to initiating slow bursts, since blocking I_h_ delayed burst onset.

Once a burst is initiated, the inward persistent sodium current (I_NaP_) can help maintain a depolarized state to prolong burst duration (Angstadt and Simone 2014; Elson and Selverston 1997; Tohidi and Nadim 2009). I_NaP_ is present in cultured spiny lobster (*Panulirus interruptus*) stomatogastric neurons, and contributes to LPG slow bursts elicited in response to synaptic input (Elson and Selverston 1997; Turrigiano et al. 1995). In *P. interruptus*, the LPG neuron is solely active in the slower gastric mill rhythm and is not electrically coupled to the pyloric pacemaker neurons (Elson and Selverston 1997; Marder and Bucher 2007). Whether a persistent sodium current contributes to LPG activity in *C. borealis* is unknown. Therefore, we asked whether the I_NaP_ blocker riluzole (30 µM) (Schuster et al. 2012; Tohidi and Nadim 2009) alters Gly^1^- SIFamide-elicited slow LPG intrinsic bursting.

LPG generated intrinsic bursts in Gly^1^-SIFamide in riluzole, but the bursts had a shorter cycle period compared to Gly^1^-SIFamide alone (Fig. 6Ai-ii). Unlike the effects of CsCl on LPG slow bursting, riluzole decreased LPG cycle period (Fig. 6B; One way RM ANOVA, F_(5,2)_ = 12.268, p = 0.004; n = 4 – 6), primarily through a decreased burst duration (Fig. 6D; One way RM ANOVA, F_(5,2)_ = 23.698, p < 0.001; n = 4 – 6), with no consistent effect on interburst interval (Fig. 6C; One way RM ANOVA, F_(5,2)_ = 1.834, p = 0.221; n = 4 – 6)., At ∼ 40 min post-riluzole, although there was some return toward control, the effects were still different from control suggesting an incomplete washout (Table 1; Fig. 6B-D). Overall, I_NaP_ was not necessary for isolated LPG slow bursting, but appeared to contribute to the active phase of this bursting in the isolated condition. This contrasts with our findings that blocking I_h_ impacted interburst interval but not burst period, suggesting that cycle period may be regulated by a balance between I_h_ and I_NaP_. However, we found a different sensitivity of slow LPG bursting to blockade of I_NaP_ when LPG was also expressing fast pyloric bursting.

**Figure 6:**
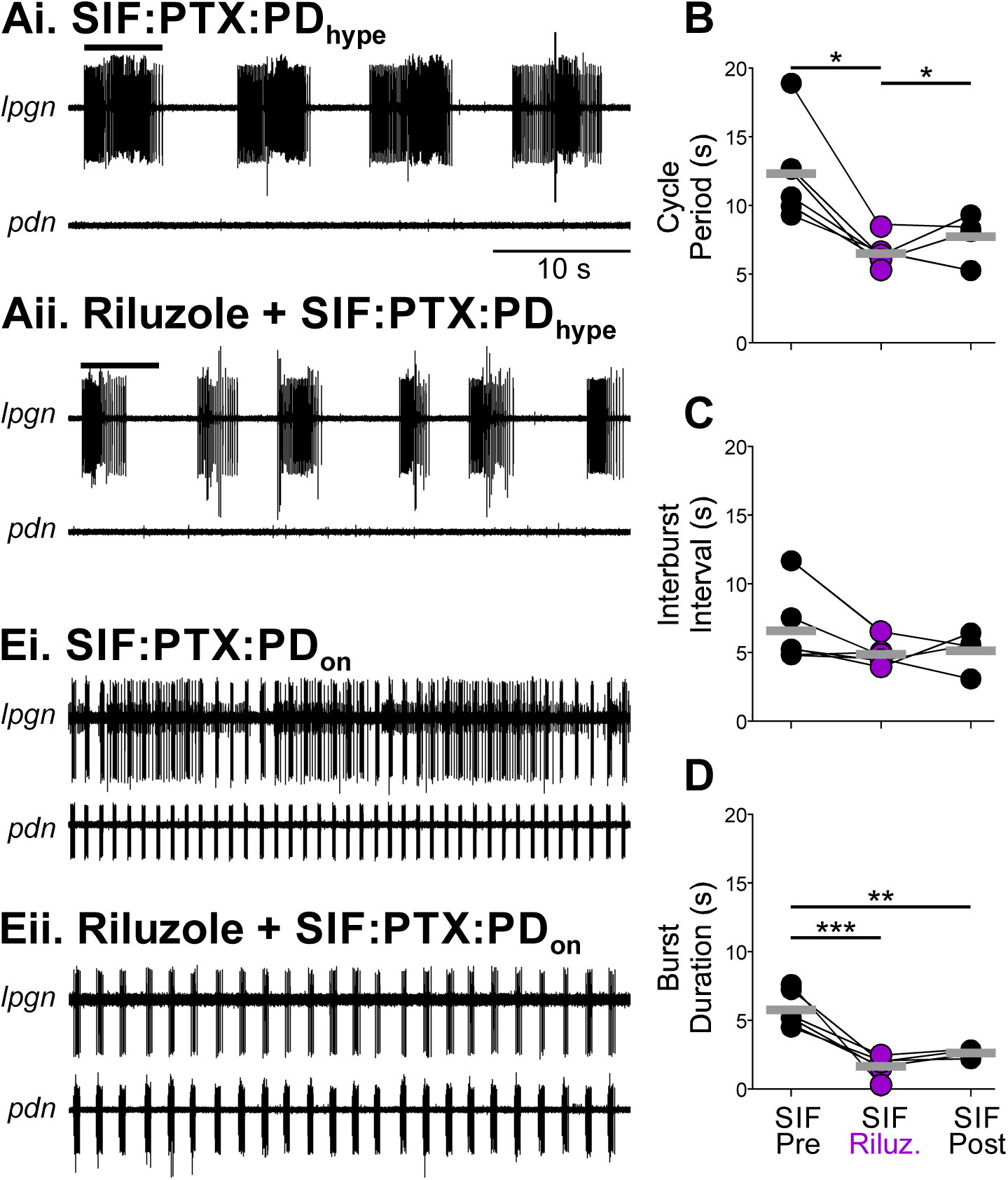
The burst duration of Gly^1^-SIFamide-elicited LPG intrinsic bursting was shorter in the presence of the I_NaP_ blocker riluzole. Ai) Gly^1^-SIFamide elicited slow intrinsic bursting in the PTX:PD_hype_ (*pdn*) isolated condition, monitored with extracellular recordings of the LPG neurons (*lpgn*). (Ai) Intracellular recording from an LPG neuron (top trace) in the presence of PTX with the PD neurons (*pdn*) held hyperpolarized (PTX:PD_hype_). 5 μM Gly^1^-SIFamide elicits intrinsic bursts during steady-state effects. Aii) In the same preparation, 5 μM Gly^1^-SIFamide with 30 μM riluzole to block I_Nap_ also elicited intrinsic bursts in LPG. Cycle period (B), burst duration (C) and interburst interval (D) in Gly^1^-SIFamide plus PTX:PD_hype_ before (SIF Pre) and after (SIF Post) Gly^1^-SIFamide plus PTX:PD_hype_ application in riluzole (SIF Riluzole); one-way RM ANOVA, Bonferroni). (**p<0.01, ***p<0.001; B-D, n = 8). Grey bars in graphs represent mean values in each condition (Gly^1^-SIFamide:PTX:PD_hype_, filled circles; Riluzole + Gly^1^-SIFamide:PTX:PD_hype_, purple circles). Ei) Application of 5 μM Gly^1^-SIFamide in the PTX:PD_on_ condition enabled periodic escapes from the pyloric rhythm and generation of several slow bursts. Eii) In Gly^1^-SIFamide in riluzole (30 μM) in the PTX:PD_on_ condition, LPG failed to escape and generate slow bursts during steady-state. Instead, LPG remained active in pyloric time (*lpgn*), coactive with the PD neurons (*pdn*). Scale bar in A applies to A and E, all recordings are from the same experiment.

When Gly^1^-SIFamide was applied in in the PTX:PD_on_ condition, LPG generated dual frequency bursts in both pyloric and gastric mill time (Fig. 6Ei-Eii). With riluzole (30 µM) added, Gly^1^-SIFamide produced variable effects on the pyloric rhythm in the PTX:PD_on_ condition. Specifically, in 3/6 preparations, the PD neurons failed to maintain rhythmic bursting in riluzole plus Gly^1^-SIFamide. In the 3/6 preparations in which fast rhythmic PD active persisted, LPG failed to escape the pyloric rhythm and generate slow bursts at steady-state in 2/3 preparations (e.g., Fig. 6Eii). In the remaining one preparation in which the pyloric rhythm remained active in SIF:PTX:PD_on_ + riluzole (30 µM), LPG generated only 8 slow bursts during a 200 s window of steady-state Gly^1^- SIFamide application. For comparison, in the control SIF:PTX:PD_on_ condition, there were 18.7 ± 5.0 bursts (n = 3) during a 200 s window of steady-state Gly^1^-SIFamide actions. As LPG only produced a few slow bursts in only one experiment, we did not quantify these burst parameters. Two additional experiments were performed in the absence of PTX in which hyperpolarizing current was used to silence the LP neuron and remove chemical synaptic inhibition to LPG (LP_hype_) as an alternative to PTX (Fahoum and Blitz 2021). In the LP_hype_ experiments, LPG produced only one slow burst (n = 1/2) or entirely failed to escape the pyloric rhythm and generate intrinsic slow bursts at steady state (n = 1/2) during riluzole plus Gly^1^-SIFamide application. In summary, in 3/5 preparations in which the pyloric rhythm remained active (2/3 PTX:PD_on_; 1/2 LP_hype_), riluzole prevented LPG from escaping rhythmic pyloric input to generate slower intrinsic bursts. In the remaining two experiments, LPG generated only 1 or 8 bursts across a 200 s window. Thus, unlike the isolated LPG neuron, in which I_NaP_ is important for burst duration, but is not essential, when LPG is receiving fast rhythmic input through electrical synapses, I_NaP_ is extremely important for burst occurrence.

Given the complementary actions of I_h_ and I_NaP_ on interburst interval and burst duration, respectively, we next asked whether co-applying their blockers was enough to eliminate intrinsic LPG bursting. In the PTX:PD_hype_ condition, application of Gly^1^- SIFamide elicited intrinsic bursting in LPG (Fig. 7Ai). When Gly^1^-SIFamide was applied in the presence of CsCl and riluzole to block both I_h_ and I_NaP_, LPG was still able to generate intrinsic bursts (n = 4) (Fig. 7Aii). The riluzole/CsCl combination caused two changes in the LPG slow bursts, an increased interburst interval and a decreased burst duration (Fig. 7), similar to the individual effects of CsCl and riluzole, respectively.

**Figure 7:**
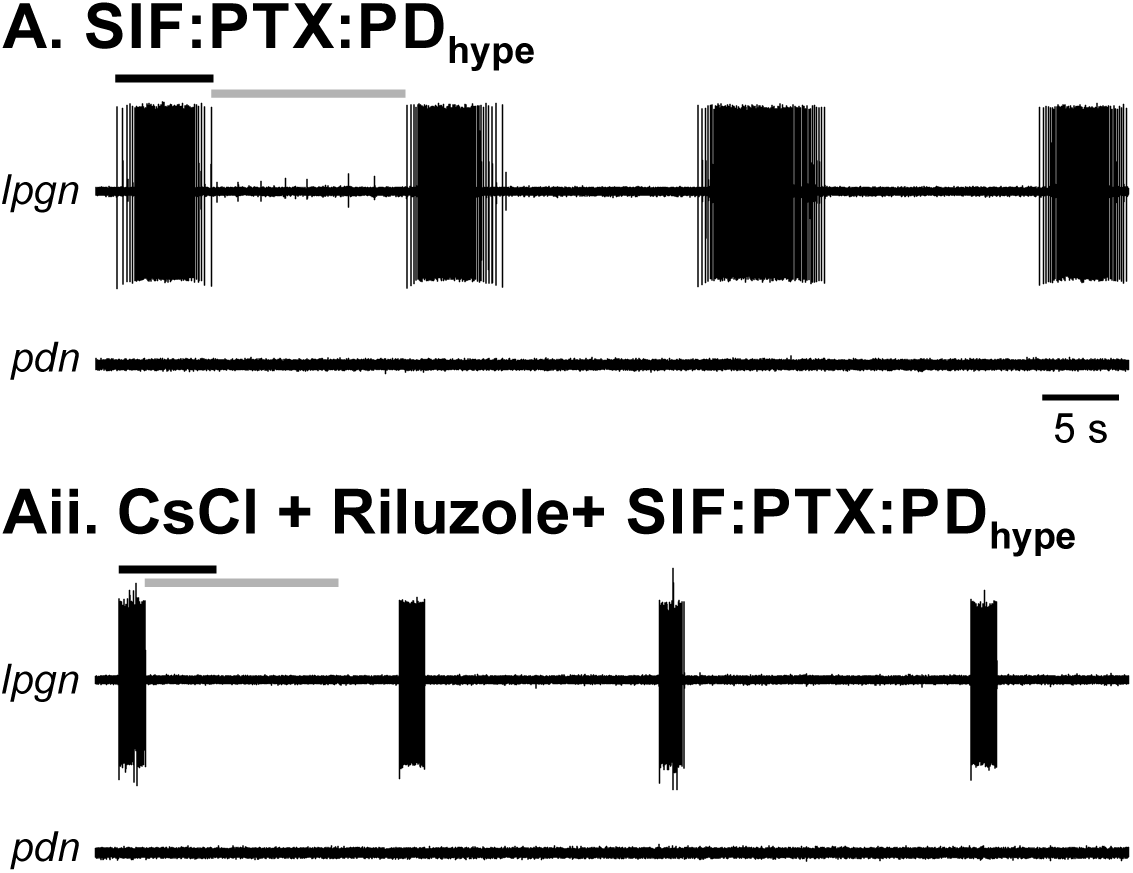
Blocking both I_h_ and I_NaP_ did not abolish LPG intrinsic bursting. Ai) In the control condition of Gly^1^-SIFamide and PTX with the PD neurons held hyperpolarized (SIF:PTX:PD_hype_) LPG generated intrinsic slow bursting (*lpgn*). PD neurons remained silent (*pdn*). Aii) In the presence of 5 μM Gly^1^-SIFamide with 30 μM riluzole and 5 mM CsCl to block I_h_ and I_Nap_, LPG generated intrinsic bursts with a shorter burst duration and longer interburst interval. Black (burst duration) and gray (interburst interval) bars from (Ai) are replicated for comparison. Scale bars apply to both panels. Recordings in are from the same preparation.

However, the persisting ability of LPG to generate slow bursts in the presence of these two blockers, indicated that an additional current(s) is sufficient for intrinsic LPG bursting, at least in the isolated SIF:PTX:PD_hype_ condition.

Although Gly^1^-SIFamide decreased the extent of SFA in LPG, there was still some SFA in the modulated condition (Fig. 3). A common mechanism for SFA is the activation of outward I_KCa_ currents during sustained activity. Thus, we next assessed the role of calcium-related currents in LPG intrinsic bursting as currents such as I_KCa_ and I_CAN_ can contribute to intrinsic bursting in many cells (Guinamard et al. 2013; Le et al. 2006; Rodriguez et al. 2013; Russell and Hartline 1982; Schneider et al. 2021). To test for a role of Ca^2+^-related currents, we used a low Ca^2+^ saline with a maintained divalent cation concentration (low Ca^2+^; 0.1x [Ca^2+^]; Mn^2+^ substitution) to decrease calcium influx. This saline decreases calcium influx sufficiently to block transmitter release and outward Ca^2+^-sensitive currents such as I_KCa_ currents in the STNS (Beenhakker et al. 2007; Blitz and Nusbaum 1999; Coleman et al. 1995; Golowasch and Marder 1992; Hooper et al. 1986; Khorkova and Golowasch 2007).

Low Ca^2+^ in the SIF:PTX:PD_hype_ condition failed to completely abolish isolated LPG slow bursting. Instead, Gly^1^-SIFamide in low Ca^2+^ saline elicited a series of progressively weaker intrinsic LPG bursts (n = 4) coincident with an increasingly depolarized trough membrane potential, eventually devolving into a semi-tonic firing at steady-state (Fig. 8A). Expanded sections of the LPG recording in figure 8A at onset of Gly^1^-SIFamide actions (Fig. 8Ai, *left*) and at steady-state (Fig. 8Ai, *right*), better illustrate the depolarized trough potential and the weakened bursting. In the same experiment, the trough voltage remained stable from onset to steady-state in the control SIF:PTX:PD_hype_ condition (Fig. 8B). Across three experiments with an intracellular LPG recording, there was a larger difference in the LPG trough membrane potential between onset and steady-state in Gly^1^-SIFamide in low Ca^2+^ compared to SIF:PTX:PD_hype_ (steady-state trough voltage – onset trough voltage in **SIF vs. lowCa:SIF**; Experiment 1: **0.0 vs. 5.1 mV**; Experiment 2: **3.4 vs. 4.2 mV**; Experiment 3: **0.0 vs. 14.0 mV**). These data suggest that one or more calcium or calcium-related currents are necessary to hyperpolarize the membrane between bursts to activate the hyperpolarization-activated current(s) required for initiating subsequent bursts, and possibly de-inactivate inward currents contributing to burst strength.

**Figure 8:**
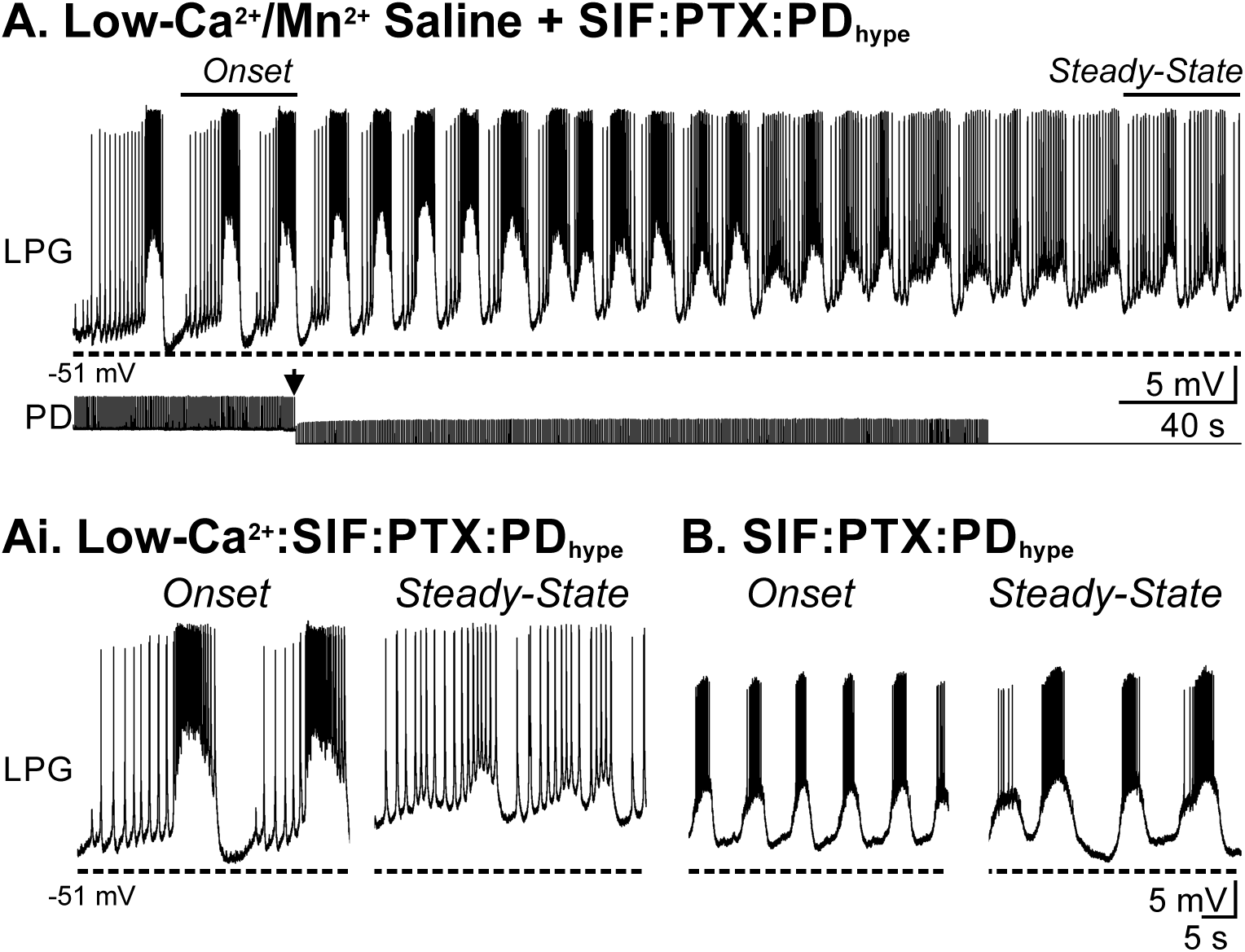
In low Ca^2+^ saline, Gly^1^-SIFamide-elicited intrinsic bursting progressively weakened, and trough voltage progressively depolarized in the isolated LPG neuron. A) Intracellular recording from an LPG neuron (top trace) in PTX with the PD neurons hyperpolarized (PTX:PD_hype_; PD neuron bottom trace) in 0.1X [Ca^2+^] saline (equimolar Mn^2+^ substitution). The dashed line indicates the initial trough membrane potential. A 400 second window demonstrates onset and development of Gly^1^-SIFamide effects across an application. The small events in PD are ectopic spikes that passively propagate to the somatic recording site. Ectopic spikes can be activated via modulation of secondary spike initiation zones along the axon (Bucher and Goaillard 2011). Gly^1^-SIFamide occasionally activates ectopic PD neuron spiking which does not alter LPG activity (Fahoum and Blitz, unpublished). At the arrow, the amount of hyperpolarizing current injected into PD was increased and the unbalanced bridge recording saturated toward the end of the time shown (straight line). Ai) Regions from A (bars and labels: *Onset* and *Steady-State*) are shown at an expanded time scale to highlight the changes in bursts between the onset and steady-state of Gly^1^-SIFamide effects in low Ca^2+^ saline. Dashed lines mark trough potential levels at onset of Gly^1^- SIFamide effects in low Ca^2+^. B) Traces from the respective control application of Gly^1^-SIFamide in PTX:PD_hype_ at onset (*left*) and during steady-state (*right*). Dashed lines indicate trough potential values at onset of Gly^1^-SIFamide effects in low Ca^2+^ PTX for comparison. Scale bars in B apply to Ai and B. All recordings are from the same preparation.

To determine whether calcium-related currents make similar contributions to LPG slow bursting in the dual-bursting condition, we assessed Gly^1^-SIFamide actions in low Ca^2+^ saline in the PTX:PD_on_ condition. In an example experiment, LPG generated a series of pyloric-timed bursts during onset of Gly^1^-SIFamide effects (Fig. 9A, Ai).

**Figure 9:**
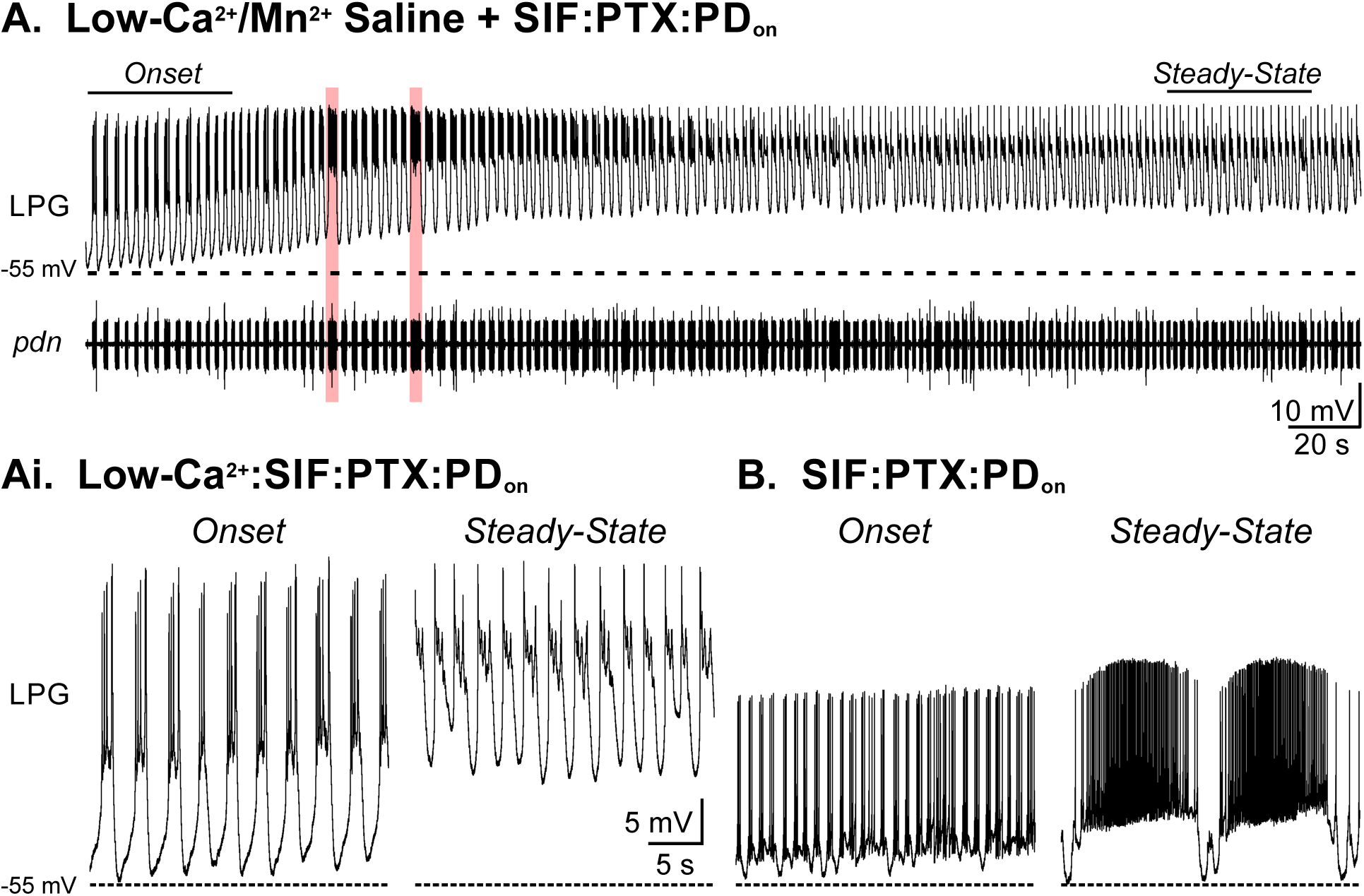
In low Ca^2+^ saline with the pyloric rhythm intact, Gly^1^-SIFamide-elicited intrinsic bursting was blocked, and trough voltage progressively depolarized. A) Intracellular recording from an LPG neuron (top trace) in the presence of PTX with the pyloric rhythm intact (PTX:PD_on_; *pdn* bottom trace) in 0.1X [Ca^2+^] saline (equimolar Mn^2+^substitution). The black dashed line indicates the initial trough membrane potential. The onset and development of Gly^1^-SIFamide effects are evident across a 400 s window. Ai) In regions from A (bars and labels: *Onset* and *Steady-State*) displayed at an expanded time scale, it is evident that LPG only produces pyloric-timed bursting and no slow bursts at onset or steady-state of Gly^1^-SIFamide effects in low Ca^2+^ saline when the pyloric rhythm is active. Note the depolarized troughs (Ai, *Steady-State*) compared to onset bursts (Ai, *Onset*) and degradation of spike amplitude. The dotted line marks the trough voltages at onset of Gly^1^-SIFamide effects in low Ca^2+^ saline. B) Expanded traces from the respective control application of Gly^1^-SIFamide in PTX:PD_on_ at onset (*left*) and during steady-state (*right*). Dotted lines indicate trough potential values at onset of Gly^1^-SIFamide effects in low Ca^2+^ PTX for comparison. Scale bars in Ai apply to Ai and B. All recordings are from the same preparation.

Throughout the application, the LPG interburst membrane potential progressively depolarized until reaching a steady-state with spike inactivation during bursts (Fig. 9A, Ai). Although there were several longer duration LPG bursts (2.4 – 2.9 s; Fig. 9A), they coincided with prolonged PD bursts (Fig. 9A, red boxes). Expanded sections of the LPG recording in figure 9A highlight the depolarization of the LPG trough voltage from onset (Fig. 9Ai, *left*) to steady-state (Fig. 9Ai, *right*) Gly^1^-SIFamide in low Ca^2+^ saline effects.

The trough membrane potential was consistently more depolarized at steady state compared to at the onset of Gly^1^-SIFamide in low Ca^2+^ saline (onset: -54.8 ± 0.9; steady-state: -44.4 ± 1.5 mV; paired *t*-test, *t*_(5)_ = -9.365, p = 0.0002, n = 6). The progressive depolarization of trough voltage did not occur in the control SIF:PTX:PD_on_ application (Fig. 9Bi) (onset: -54.3 ± 1.5; steady-state: -52.1 ± 1.5 mV; paired *t*-test, *t*_(5)_ = -1.919, p = 0.113, n = 6). The depolarized trough voltage during Gly^1^-SIFamide in low Ca^2+^ saline was similar to that occurring in the isolated LPG bursting in the low Ca^2+^ SIF:PTX:PD_hype_ condition (Fig. 8). However, there was a greater impact of low Ca^2+^saline on LPG slow bursting when the faster pyloric oscillations were present.

Specifically, in 7/11 experiments where rhythmic PD activity persisted in low Ca^2+^ saline plus Gly^1^-SIFamide, LPG failed to escape from the faster pyloric input and produce any slow, intrinsic bursts. In the other 4/11 experiments, LPG escaped the pyloric rhythm to produce 1-3 slow, intrinsic bursts during the onset of Gly^1^-SIFamide effects in low Ca^2+^ saline before bursting stopped and LPG became either silent or tonic during steady-state. Together, these findings suggest that one or more calcium or calcium-related currents are necessary for periodic escapes from the pyloric rhythm, and further support that calcium-related current(s) are necessary for burst termination and hyperpolarization between bursts.

## Discussion

Previously, we identified a modulatory input and its neuropeptide transmitter that switch a neuron from fast only to dual fast/slow frequency oscillations (Blitz et al. 2019; Fahoum and Blitz 2021). While the fast oscillations are due to electrical coupling among the fast network pacemaker neurons, the slow oscillations are due to intrinsic properties of the switching neuron and are not dependent upon synaptic input from the second network (Fahoum and Blitz 2021). In this study, we identified multiple intrinsic properties that are modulated by the neuropeptide Gly^1^-SIFamide, and several candidate ionic currents that could contribute to the intrinsic bursting. Furthermore, the sensitivity of LPG slow bursting to blocking currents differed when isolated from both networks versus the dual-frequency condition. This suggests important interactions between electrical coupling input and intrinsic currents in a switching neuron.

### Neuronal Switching

Neuromodulators can induce neurons to alter their network participation in invertebrate and vertebrate neural networks (Bartlett and Leiter 2012; Bouret and Sara 2005; Dickinson 1995; Dickinson et al. 1990; Nagy and Dickinson 1983; Oyarzabal et al. 2022; Tryba et al. 2008). Neuronal switching includes a neuron changing participation from one network to another, switching between single- and dual-network participation, or reconfiguration of neurons from multiple networks into a novel network (Bouret and Sara 2005; Dickinson et al. 1990; Meyrand et al. 1994; Weimann et al. 1991; Weimann and Marder 1994). Thus far, the mechanisms by which a neuromodulator elicits neuronal switching are mostly identified in smaller invertebrate networks. In general, neuromodulators alter synaptic and/or intrinsic membrane properties of network neurons (Harris-Warrick 2011; Marder and Thirumalai 2002; Nadim and Bucher 2014). However, the most well-established cellular mechanism for a neuromodulator to elicit a switch in participation is modification of synaptic strength (Hooper and Moulins 1990; Meyrand et al. 1994; Weimann and Marder 1994). More recent studies highlight an alternate possibility where modulation of intrinsic properties can underlie neuronal participation (Drion et al. 2019; Fahoum and Blitz 2021; Tryba et al. 2008). For example, in large mammalian respiratory networks, modulation of intrinsic properties releases neurons from dual-network participation, although modulation of electrical synapses remains a potential mechanism (Tryba et al. 2008). Further, in the small STNS feeding networks, we found that neuronally-released, or bath-applied neuropeptide Gly^1^- SIFamide enables oscillations at a second frequency selectively through modulation of intrinsic properties (Fahoum and Blitz 2021). However, the ion channel targets and intrinsic membrane properties that are altered by this modulatory pathway were not identified. In a computational study, modulation of a slow conductance determined whether a hub neuron participated in single- versus dual-network activity, but these findings were not tested in a biological system (Drion et al. 2019).

### Modulation of Intrinsic Properties

Intrinsic oscillations occur due to a combination of voltage-gated ionic currents contributing to different components of the oscillation. The ability to generate intrinsic oscillations can be enabled or disabled by neuromodulators which alter these ion channel properties (Adams and Benson 1985; Harris-Warrick 2010; Marder and Calabrese 1996; Peña et al. 2004). Different ion channels, or combinations of channels, underlie other intrinsic properties within neurons which may contribute to intrinsic oscillations, or further shape neuronal activity. For instance, in many CPG and other rhythmic networks, inhibition can activate, or de-inactivate voltage-gated currents which in turn drive the active phase of bursting (Bucher et al. 2015; Lu et al. 2020; Marder and Calabrese 1996). Activation of the I_h_ current can underlie this drive toward burst threshold from a hyperpolarized membrane potential. The I_h_ current typically elicits a depolarizing sag in membrane potential in response to sustained inhibition and can produce a PIR above baseline following removal of the inhibition (Angstadt et al. 2005; Goaillard et al. 2010; Hogan and Poroli 2008; Sangrey and Jaeger 2010; Yang et al. 2018). Here we found that LPG expresses a small amount of PIR in control conditions which is enhanced by Gly^1^-SIFamide. This was accompanied by an increased sag potential in Gly^1^-SIFamide, suggesting modulation of I_h_, which is a target of several neuromodulators in the STNS (Ballo et al. 2010; Krenz et al. 2015; Peck et al. 2006). Furthermore, in the presence of the I_h_ blocker CsCl, the interburst interval of Gly^1^- SIFamide-elicited LPG slow bursts was extended with no consistent effect on burst duration. Similar contributions of I_h_ to interburst interval occur in the lobster pyloric network (dopamine modulation) and leech heartbeat system (myomodulin peptide modulation) (Peck et al. 2006; Tobin and Calabrese 2005). In the LPG neuron, it appears that I_h_ can speed up the rate of depolarization to the next burst, but it is not necessary for slow intrinsic bursting.

In many bursting neurons, a voltage-dependent, persistent current contributes to maintaining activity during the burst. In particular, a non-inactivating modulator activated, voltage-dependent, mixed cation inward current (I_MI_) is an important contributor to fast (pyloric) and slow (gastric mill) bursting in STG neurons (DeLong et al. 2009; Golowasch et al. 2017; Gray et al. 2017; Kintos et al. 2016; Rodriguez et al. 2013; Swensen and Marder 2000). Similarly, in chewing, respiratory, and locomotor network neurons in rodents and leeches, a persistent Na^+^ current supports bursting (Crisp et al. 2012; Harris-Warrick 2010; Tazerart et al. 2008; Yamanishi et al. 2018). The I_MI_ current is activated by many peptides in the STNS, however there is no effective blocker for this current (Gray et al. 2017; Schneider et al. 2021). We did not perform voltage clamp analyses in this study and therefore did not determine if I_MI_, or the newly identified transient I_MI_ (I_MI-T_) (Schneider et al. 2021) is a target of Gly^1^-SIFamide.

However, we tested whether the I_NaP_ blocker riluzole impacted Gly^1^-SIFamide-elicited LPG bursting. We found that slow intrinsic LPG bursting was still elicited but the duration of Gly^1^-SIFamide-elicited bursts was decreased in riluzole, without an effect on the interburst interval. This suggests that I_NaP_ is important for burst maintenance but not burst onset. I_NaP_ is present in the LPG neuron in lobster and in unidentified cultured lobster STG neurons (Elson and Selverston 1997; Turrigiano et al. 1995). Although I_h_ and I_NaP_ have complementary roles (burst initiation and maintenance, respectively), applying both CsCl and rilzuole was not sufficient to block Gly^1^-SIFamide-elicited bursting. This may be due to incomplete block, or Gly^1^-SIFamide modulation of an additional current such as I_MI_.

Gly^1^-SIFamide does modulate additional intrinsic properties and thus, likely additional currents. In particular, this peptide decreased the extent of SFA in LPG. SFA can contribute to intraburst firing frequency and burst duration, and often plays an important role in regulating burst termination (Ha and Cheong 2017; Katz and Frost 1997; El Manira et al. 1994). There was also a general increase in LPG excitability, measured as a larger F/I ratio, which may be due to the same or additional mechanisms as those underlying the decreased SFA. The increased excitability and enhanced firing during Gly^1^-SIFamide application may contribute to the ability of LPG to maintain a burst even when its electrically coupled partners AB and PD terminate their pyloric-timed bursts. Calcium activated K^+^ current(s) (I_KCa_) often contribute to SFA and their modulation can regulate bursting (Ha and Cheong 2017; Harris-Warrick 2010; El Manira et al. 1994; Overton and Clark 1997; Ramirez et al. 2012). SFA was decreased, but not eliminated in Gly^1^-SIFamide and thus may be a contributing factor in burst termination. A role for the outward I_KCa_ is supported by the progressive depolarization of the LPG trough that we found during Gly^1^-SIFamide application in low Ca^2+^ saline.

In addition to a progressive depolarization of the LPG trough, LPG slow bursting was greatly reduced in the isolated condition or eliminated entirely in the intact pyloric network condition when Gly^1^-SIFamide was applied in low Ca^2+^ saline. This further supports a role of I_KCa_. If there is not sufficient calcium influx to activate I_KCa_, the membrane potential is not sufficiently hyperpolarized to activate I_h_, or to de-inactivate other inward currents and there is no, or weakened, drive to initiate a new burst.

However, it is also possible that a calcium current more directly contributes to the LPG slow bursting. One possibility for this is the transient, low-threshold calcium-dependent current (I_Trans-LTS_). The slow activation of I_Trans-LTS_ can provide a depolarizing influence that, when combined with another inward current such as I_h_, is sufficient to initiate bursts in the gastric mill neuron LG in the crab STNS on a similar time scale as LPG slow-bursting (Rodriguez et al. 2013). Further, the transient nature of I_Trans-LTS_ and its ability to recruit outward currents may contribute to burst termination (Rodriguez et al. 2013; Schneider et al. 2021). Involvement of the calcium-dependent I_Trans-LTS_ would be consistent with the decline in burst quality we found in low Ca^2+^ saline. A possible dual contribution of I_h_ and I_Trans-LTS_ to PIR in LPG may also explain why cesium blockade of I_h_ extended the interburst interval but did not block intrinsic bursting.

### Co-modulation of multiple properties

A balance of inward and outward currents is necessary for intrinsic bursting (Amarillo et al. 2014; Ellingson et al. 2021; Golowasch et al. 2017). Thus, it is not surprising that Gly^1^-SIFamide would modulate more than one intrinsic property and thus more than one intrinsic current. Correlations occur between ion channels at the mRNA and conductance levels in multiple cell types (Khorkova and Golowasch 2007; Schulz et al. 2006; Tran et al. 2019; Yang et al. 2022). Co-modulation of multiple intrinsic properties, which can maintain such correlations, appears to be important for increasing the range over which rhythmic networks produce stable oscillations (Ellingson et al. 2021). For the LPG neuron, and other dual-frequency oscillators, it is not only a balance of multiple intrinsic currents, but also a balance between intrinsic and synaptic currents that is likely necessary to enable oscillations at both frequencies.

In the intact dual-network condition, LPG receives fast rhythmic electrical coupling input from the AB/PD neurons (Marder and Eisen 1984; Shruti et al. 2014). We found differential importance of I_NaP_ and calcium-related currents in the isolated LPG versus intact network conditions. The inability of LPG to periodically escape the rhythmic electrical input in the presence of riluzole suggests that the electrical synaptic input can dominate LPG activity without an additional inward current such as a persistent sodium current. This could either be due directly to the inward current, or due to the additional leak decreasing the efficacy of the electrical coupling current during LPG slow bursts.

It is not yet known if Gly^1^-SIFamide modulates gap junctions. This is a possibility as electrical synapses can be modulated by both amines and peptides (Lane et al. 2018; Zsiros and Maccaferri 2008). In snail neurons, simulated addition of electrical synapses through dynamic clamp supported bistability and dual network activity (Bem et al. 2005). Additionally, changes in ionic currents and input resistance can alter coupling between cells (Alcamí and Pereda 2019; Nadim et al. 2017; O’Brien 2014). This may explain why LPG could not periodically escape the pyloric rhythm when I_NaP_ was blocked. If this is the case, it is possible that Gly^1^-SIFamide may regulate electrical synapses through multiple pathways – both directly and indirectly through modulation of other currents. Regardless of the mechanism, changes in functional coupling strength may be necessary for enabling slow bursts.

### Functional Implications

Network switching extends the functional range of networks and enables changes in the coordination between networks (Bouret and Sara 2005; Dickinson 1995; Meyrand et al. 1994; Tryba et al. 2008; Weimann and Marder 1994). Neuromodulation is suggested to elicit rapid rhythmic network reconfiguration among larger circuits as well. For example, norepinephrine appears to contribute to rapid changes in synchronization of multiple vertebrate brain regions during switches in attention (Bouret and Sara 2005; Oyarzabal et al. 2022; Ramirez et al. 2016). Disordered inter-network coordination and decreased flexibility in oscillatory networks can lead to dysphagia, memory and locomotor deficits, and disruption of cognitive processes (Barlow 2009; Moore et al. 2014; Rangel et al. 2016). Our findings highlight the importance of considering modulation of intrinsic properties (Drion et al. 2019; Tryba et al. 2008), either alone, or in combination with modulation of synaptic properties, when investigating mechanisms underlying flexibility of network participation among related networks in other systems.

## ENDNOTE

At the request of the author(s), readers are herein alerted to the fact that additional materials related to this manuscript may be found at https://github.com/blitzdm/Snyder_Blitz_2022. These materials are not a part of this manuscript and have not undergone peer review by the American Physiological Society (APS). APS and the journal editors take no responsibility for these materials, for the website address, or for any links to or from it.’

## Acknowledgements

We thank Savanna-Rae Fahoum and Barathan Gnanabharathi for reading earlier versions and providing helpful discussions, Michael Hughes and the Statistical Consulting Center at Miami University for assistance with statistical analysis, and Kathleen Killian and Paul James for helpful comments. This work funded by National Science Foundation (IOS: 1755283; DMB) and the Biology Department at Miami University.

